# Lipid Dysregulation Unveil the Intricate Interplay of Lysosomal and Mitochondrial Changes in Frontotemporal Dementia with GRN Haploinsufficiency

**DOI:** 10.1101/2024.01.22.576606

**Authors:** Jon Ondaro, Jose Luis Zúñiga-Elizari, Mónica Zufiría, Maddi Garciandia-Arcelus, Miren Zulaica, Miguel Lafarga, Javier Riancho, Ian James Holt, Adolfo López de Munaín, Fermin Moreno, Francisco Javier Gil-Bea, Gorka Gereñu

**Affiliations:** 1Department of Neuroscience, Biodonostia Health Research Institute (IIS Biodonostia), San Sebastian, Spain; Center for Biomedical Research of Neurodegenerative Diseases (CIBERNED), Madrid, Spain; Department of Anatomy and Cell Biology, University of Cantabria-IDIVAL, Santander, Spain; Hospital General Sierrallana-IDIVAL, Torrelavega, Spain; Department of Medicine and Psychiatry. University of Cantabria, Santander, Spain; Department of Clinical and Movement Neurosciences, Queen Square Institute of Neurology, Faculty of Brain Sciences, University College London, London, United Kingdom; Donostia University Hospital, San Sebastian, Spain; IKERBASQUE Basque Foundation for Science, Bilbao, Spain; Faculty of Health Sciences, Public University of Navarra, Pamplona, Spain; Department of Physiology, Faculty of Medicine and Nursing, University of Basque Country (UPV-EHU), Leioa, Spain

**Keywords:** Lysosome, Mitochondria, Granulin haploinsufficiency, Frontotemporal dementia, Lipid metabolism, Neurodegeneration

## Abstract

This study investigates the cellular pathology resulting from haploinsufficiency of progranulin (PGRN) in frontotemporal dementia (FTD) associated with granulin (GRN) mutations. Utilizing fibroblasts from FTD patients carrying a distinctive GRN mutation (c.709-1G>A), we observed lysosomal and lipofuscin accumulation, impaired lysosomal function, compromised autophagic flux, and mitochondrial abnormalities. Notably, recombinant human progranulin (rhPGRN) treatment restored lysosomal acidification, mitigated mitochondrial defects, and demonstrated beneficial effects. FTD-GRN fibroblasts exhibited abnormal lipid metabolism with increased lipid droplet formation, influenced by GRN haploinsufficiency and modulated by rhPGRN. Under nutrient-rich conditions, lipid droplet dynamics were shaped by autophagy and mitochondrial processes, potentially due to impaired fatty acid oxidation. These findings highlight a direct association between GRN deficiency and altered lysosomal-mitochondrial interactions, influencing lipid metabolism and contributing to FTD pathogenesis. The documented lysosomal dysfunction, impaired autophagy, mitochondrial anomalies, and altered lipid metabolism collectively suggest a complex interplay of cellular processes in the development of FTD-GRN.

## INTRODUCTION

Frontotemporal dementia (FTD) is a neurodegenerative disorder and the second leading cause of dementia, affecting patients in behavior and language capabilities under 65 years old [11, 23]. Despite ongoing progress, there are no effective treatments for FTD. A majority of FTD cases are sporadic, being considered aging the most prominent risk factor. However, 40% of patients have a family history of dementia, suggesting a substantial genetic involvement in FTD pathogenesis [68]. Mutations in multiple genes such as microtubule associated protein tau (*MAPT*), granulin (*GRN*), FUS RNA binding protein (*FUS*), valosin containing protein (*VCP*) and C9orf72-SMCR8 complex subunit (*C9orf72*) are considered causative factors to develop Familial FTD (fFTD). Among them, mutations in GRN gene account for 5-20% of the total fFTD cases [65]. These mutations lead to an haploinsufficiency of its coding protein progranulin (PGRN), resulting in a PGRN truncated protein production [31] and triggering neurodegeneration. One of these FTD-causing mutations was discovered 15 years ago in a cluster of Basque families carrying an ancestral distinctive genetic alteration (c.709-1G>A) in the GRN gene [47].

FTD-GRN is characterized by TAR DNA-binding protein 43 (TDP-43) pathology, developing TDP-43 cytoplasmic inclusions in the brain [48]. Additionally, FTD-GRN brain exhibits other specific neuropathological features, including the accumulation of lipofuscin granules [79], and abnormally digested lysosomes [54]. These accumulations are particularly present in the brains of patients with Neuronal Ceroid Lipofuscinosis (NCL), a lysosomal storage disease (LSD) that can be caused by specific GRN mutations that promote a complete absence of PGRN [70].

PGRN is a secreted glycoprotein composed of 7.5 granulin segments (granulin A-G, and paragranulin). It plays a significant role in numerous cellular processes [55]. While the precise function of PGRN remains not fully understood, individual granulins can fulfill diverse and even opposing functions compared to the full-length PGRN. Importantly, increasing evidence has highlighted the pivotal role of PGRN in lysosomal function [12, 19, 38, 62]. Intracellularly, full-length PGRN is predominantly localized within the lysosome, and is proteolytic cleavage by lysosomal proteases [36, 42, 53], leading to the production of individual granulin peptides. These peptides are believed to be active to regulate lysosomal enzymes, such as cathepsin D [14, 17, 76, 83] or glucocerebrosidase (GCase) [6, 46, 77, 84]. Further, the *GRN* gene contains two potential coordinated lysosomal expression and regulation (CLEAR) sequences to interact with transcription factor EB (TFEB). TFEB controls the expression of lysosomal genes in response to lysosomal requirements arising from increased nutrient demands during starvation (STV)-induced Serine/threonine**-**protein kinase mTOR (mTOR) inhibition [50, 72]. Consequently, lysosomal dysfunction can play a crucial role in FTD-GRN neurodegeneration [63].

The lysosome is a dynamic organelle that, through its role in cellular waste and recycling, is proficient in centrally regulating nutrient sensing and reconfiguring metabolism of the cell in response to energy demand, to maintain cellular homeostasis and ensure survival [61]. To fulfill this role, the lysosome interacts with other cellular structures exchanging content and information through establishing membrane contact sites [10]. For instance, the lysosome exerts a significant mechanism of metabolic control by facilitating the quality control of mitochondria through mitophagy, which involves the selective removal of dysfunctional mitochondria. Hence, as mitochondria are high-demanding organelles, especially in non-dividing cells like neurons, they heavily rely on lysosomal mitophagy to uphold cellular energy homeostasis, rendering it an indispensable aspect of metabolic regulation [7].

The ability of the cells to adapt and reconfigure nutrient utilization is a key factor in maintenance of metabolic balance during energy scarcity. This reconfiguration involves a shift away from glycolysis towards mitochondrial β-oxidation of fatty acids (FAs) [26] that originate from stored triacylglycerol (TAG) or cholesterol esters within lipid droplets (LDs). During energy stress, macro-autophagy is enhanced to recycle cellular components and supply the cell with essential constituents to prioritize vital biological processes [28, 81]. Within the lysosome, membrane lipids are broken down into FAs and released. These FAs can serve as a source of energy production for mitochondria [18, 35, 66] or be re-esterified and packaged into new LDs in the endoplasmic reticulum, leading to a paradoxical increase in LDs [66]. This mechanism aims to cope with excessive FA influx into mitochondria and prevent lipotoxicity [8, 29, 45, 64, 78].

Therefore, alterations at the lysosomal level have the potential to impact cellular homeostasis during periods of energy demand at least at two different levels: first, by impeding the availability of FAs as an alternative fuel source, and second, by increasing the risk of accumulating aberrant metabolic organelles such as mitochondria [61]. The latter may result in less flexibility to reconfigure alternative sources of energy production. A common feature shared by mitochondria and lysosomes is that alterations in the enzymes of these organelles often lead to neurological pathologies [4, 16, 25, 32, 44], suggesting functional connections between mitochondria and lysosomes.

The aim of this study is to test the hypothesis that PGRN haploinsufficiency associated with FTD-GRN, may lead to an alteration in the crosstalk between the lysosome and the mitochondria, resulting in detrimental consequences at the metabolic level. To achieve this, we utilized human fibroblasts obtained from FTD-GRN patients, which exhibit relevant pathological features observed in FTD-GRN brains.

## MATERIALS AND METHODS

### Human samples

A total of 6 skin biopsies were analyzed: 3 from carriers of the c.709-1G>A GRN gene mutation and 3 control individuals without mutation in GRN nor any sign of neurological degeneration (Table 1). Control individuals were healthy relatives of patients. All patients were treated in the Donostia University Hospital by applying consensus criteria as published elsewhere [51. This study was approved by the Donostia University Hospital Ethical Research Board (approval no. B17-GRN-2014-01) and was conducted in accordance with the Declaration of Helsinki’s ethical standards.]. Skin biopsy and genetic analyses were performed following written, informed consent, according to the rules of the ethical committee of our Institutions.

**Table 1:** Characteristics of individuals enrolled in this study.

| Characteristic | CTL (n=3) | GRN (n=3) |
| --- | --- | --- |
| Age at biopsy | 63.0 ± 13 | 67.9 ± 6 |
| Age at onset | - | 63.0 ± 5 |
| Gender, female:male | 2:1 | 0:3 |
| Genotype | Negative | c.709-1G>A |
| Phenotype, Asymptomatic:FTD | 3:0 | 1:2 |
| Control (CTL): Individuals without sign of neurological degeneration. Patients with mutations in GRN gene (GRN); c.709-1G>A, Progranulin mutation; FTD, Frontotemporal Dementia; n, number of subjects. Values are expressed as mean ± SD. |  |  |

### Cell culture conditions

Primary fibroblast cultures were established from skin biopsy of healthy donors and FTD patients. Fibroblast cells were cultured in Cell Medium (CM), DMEM/high glucose (Gibco) with FBS (10%)(Gibco), 1.2% glutaMAX (ThermoFisher scientific) and penicillin/streptomycin (100 mg/ml) (Gibco) incubated at 37 °C and 5% CO2. To study in nutrient deprivation condition we used Starving medium, EBSS with Mg, Ca and PhenolRed (Gibco), FeCl3.6H2O (0.07 mg/mL), D-Glucose (4.5 g/L) (ITW reagents), 1 mM Piruvate, 1x Vitamin Solution, penicillin/streptomycin (100 mg/ml) (Gibco). Control SH-SY5Y cells were cultured in DMEM/F12 with FBS (10%) at 37 °C and 5% CO2. Silenced TDP43 or Control cells (shTDP43 or ShRNA SHSY-5Y cells) were generated by treating parental SH-SY5Y cells with lentiviral particles with short hairpin RNA (ShRNA), TDP43-specific shRNA lentiviral clone (clone ID TRCN000016038, Sigma Aldrich) or SHC001 (shRNA Empty Vector Control Plasmid DNA). The lentiviral particles were produced and titulated by the viral vectors unit platform of CNIC (Madrid). 50% confluent cells were treated with a MOI of 10 viral particles per cell for 24 h. Treatments: 30 µM Chloroquine (CLQ) (Sigma-Aldrich) for 5h or 100 nM Bafilomycin (BafA1) (Sellectchem) treatment or vehicle for 2h. Starving medium (FBS and amino acids free) FeCL3.6H2O [0,0669mg/L], D-Glucose [4,5 g/L], Pyruvate [1mM] (Gibco), MEM Vitamin Solution (100X), penicillin/streptomycin (100 mg/ml) (Gibco) in EBSS (calcium, magnesium, phenol red) (Gibco) for 6 h was used for high proteolysis/pro autophagic (starvation) conditions. 500ng/mL or 1000ng/mL human recombinant PGRN (rhPGRN) (R&D) for 2, 6, 12 or 24h.

### mRNA expression/processing analysis

Total RNA was extracted using the RNeasy Mini Kit with RNase-free DNase set (Qiagen). Reverse transcription was performed using the SuperScript VILO cDNA Synthesis Kit (Thermo Fisher) according to the manufacturer’s guidelines.

Quantitative real-time PCR was performed in an CFX384 Touch Real-Time PCR Detection System (Bio-Rad) using Power SYBR Green Master Mix (Thermo Fisher), 300 nM of primer pair and 10 ng of cDNA. GAPDH was used as a housekeeping gene. 2-ΔΔCt method was used for relative quantification. The primer sequences were: GRN promoter forward 5′ CTCTCCAAGGAGAACGCTACCA 3′; reverse 5′ GACTGTAGACGGCAGCAGGTAT 3′; LAMP2 forward 5′ TGACGACAACTTCCTTGTGC 3′; reverse 5′ AGCATGATGGTGCTTCAGAC 3′; and GAPDH forward 5′ ACAGTTGCCATGTAGACC 3′; reverse 5′ TTGAGCACGGGTACTTTA 3′.

The reverse transcriptase was performed with the SuperScript Vilo kit (Thermo Fisher) and the PCR was done with the ImmoMixTM reaction-mix (Meridian Bioscience) with the primer sequence PFKP promoter forward 5′ CATGGTGGACGGAGGCTC 3′; reverse 5′ CAGCCCACTCCACTCCTTC 3′. The samples were loaded on a 1% agarose gel, and the signal was visualized using an iBright FL1000 Imaging System (Thermo Fisher).

### Sample preparation and western blot

Primary fibroblast were lysed in Cell Lysis buffer (#9803, Cell Signaling) containing phosphatase and protease inhibitors (Thermo Fisher Scientific). Homogenates were centrifuged at 13000× g for 15 min 4 °. All samples were mixed with an equal volume of loading buffer (0.16 M Tris-HCl pH 6.8, 4% SDS, 20% glycerol, 0.01% bromophenol blue, 0.1 M DTT) and ran in Mini-PROTEAN® TGX™ Precast Protein Gels, Criterion™ TGX™ Precast Protein Gels, (Bio-Rad) or Tris-HCl/Tris-tricine gels. Proteins were transferred to Amersham™ Protran® Western blotting membranes, nitrocellulose (pore size 0.2 μm, roll W × L 300 mm × 4 m, pkg of 1 ea) (GE healthcare) overnight 4 °C at 15 V. After 1 h blocking in 5% BSA in TBS-Tween, Membrane was incubated with specific primary antibodies overnight 4 °C. Primary Antibodies: PGRN (1μg/mL, AF2420, R&D systems), LAMP-1(1:10 000, ab25630, Abcam), LAMP-2 (1:1 000, #49067, Cell Signaling), LC3Bxp (1:1 000, #3868, Cell Signaling), SQSTM1/P62 (1:1 000, #5114, Cell Signaling), β-Tubulin (1:5 000, MA5-16308, Thermo Fisher) PINK1 (1:1 000, BC100-494, Novus Bio) TOM20 (1:1 000, 11802-1-AP, Proteintech), GAPDH (1:50 000, 60004-1-Ig, Proteintech), Lamin A+C (1:10 000, ab108595, Abcam), TDP-43 (1:1 000, ab104223, Abcam). Either Anti-mouse/Anti-rabbit IgG, HRP-linked Antibody (1:5 000, 7076, 7074, Cell Signaling) or fluorescent secondary antibodies were used at 1/5 000, 2 h, room temperature (RT). Bands were scanned iBright FL1000 Imaging System (Thermo Fisher). The resulting images were quantified by Image Studio Lite software (LI-COR Bioscience). Immunoreactivity was calculated as the increase over the signal obtained in samples from control cells. Quantifications were normalized against loading controls (β-Tubulin).

Cell fractionation was performed by using a cell fractionation kit Abcam, ab109719) according to the manufacturer’s instructions. Nuclear–cytoplasmic fractionation was conducted using the NE-PER Nuclear and Cytoplasmic Extraction Reagents kit (Thermo Fisher Scientific) according to the manufacturer’s protocol.

### Immunofluorescence

Primary fibroblast were cultured in Ibidi μ-Slides or multi well plates (µ-Plate 96 Well Black, 89626, ibidi), when appropriate confluence was reached they were fixed in 4% paraformaldehyde for 20 min followed by blocking and permeabilization (5% BSA and 0.1% TritonX-100 in PBS) for 1h at RT. As primary antibodies, we used Anti-LC3B (#2775, Cell Signaling), LAMP-1 (ab25630, Abcam), TOM20 (11802-1-AP, Proteintech) incubated overnight at 4°C. Thereafter, cells were washed three times and incubated for 2h with secondary antibodies AlexaFluor488 or AlexaFluor647 (Life Technologies) and chromatin stained with DAPI (1μg/mL, D1306, Thermo Scientific). Primary antibody was omitted as a negative control. Samples were mounted with Ibidi mounting medium and image taken using a Zeiss LSM 900 Confocal Microscope, with time-lapse acquisition system live cell acquisition system with temperature and atmosphere control. Image analysis was performed using Image J. (Colocalization analysis). For mitochondria network analysis, the “Skeletonize 2D/3D” command was applied to the threshold images. With the “Analyze Skeleton” command we calculate the number of branches, branch length and branch junctions in the skeletonized network.

To quantify the fluorescence intensity of LC3 puncta and LAMP-22, the mean fluorescence intensity in LC3 puncta or LAMP-2 channel was calculated across the entire image. To determine the % of Optical Density, the mean fluorescence intensity was normalized with the image area.

To quantify the fluorescence intensity of LC3 in Mitochondria, we made a mask using the LC3 puncta channel, and then overlap with the TOM20 channel mask. The LC3 particles Area in Mitochondria were quantified with the ImageJ ‘‘analyze particles’’ function in threshold images, with size (square μm) settings from 0.1 to 100 and circularity from 0 to 1. To determine LC3 % area per cell, LC3 particles in Mitochondrial area was normalized with Cell area calculated with Cellpose generalist algorithm for cell segmentation.

Measurements of mitochondrial area were generated using the ‘analyze particles’ function in Fiji (NIH) with a minimum area of 0.25 mm. Measures of mitochondrial length, junctions (voxels with three or more neighbors) and branches (slab segments connecting end points to either junctions or other endpoints) were determined using the ‘skeletonize’ and ‘analyze skeleton’ plugins in Fiji (NIH). Analyses were carried out on whole cells.

### Live Cell Imaging

For the lysosomal functionality assay fibroblast cells were incubated with CM containing 100 nM LysoTracker™ Red DND-99 (Thermo Fisher) for 5h. Cells were then washed three times with CM and immediately images were taken.

For the FA Pulse and Chase Assay fibroblast cells were incubated with CM containing 1 mM BODIPY 558/568 C12 (Red C12, Life Technologies) for 16 hr. Cells were then washed three times with CM, incubated for 1 hr in order to allow the fluorescent lipids to incorporate into LDs, and then chased for the time indicated in CM or STV medium in the absence or presence of rhPGRN. LDs were labeled with 200ng/mL BODIPY 493/503 (Life Technologies) immediately prior to imaging and was present during imaging.

For FA tracing into mitochondria, Fibroblast cells were incubated with CM containing 1 mM BODIPY 558/568 C12 (Red C12, Life Technologies) for 16 hr. Cells were then washed three times with CM, incubated for 1 hr in order to allow the fluorescent lipids to incorporate, and then chased. Mitochondria were labeled with 100nM MitoTracker™ Green FM (M7514, InvitrogenTM) for 30min. Cells were then washed once with CM and immediately images were taken.

All images were acquired on a Zeiss LSM 900 Confocal Microscope, with time-lapse acquisition system live cell acquisition system with temperature and atmosphere control.

### Image Processing, Analysis, and Statistics

Images were analyzed using ZEN Blue Imaging Software (Zeiss) and ImageJ (NIH). Image brightness and contrast were adjusted in ImageJ (NIH). To quantify the fluorescence intensity of Red C12 in LDs, we made a mask using the BODIPY 493/503 channel, and then overlap with the Red C12 channel mask. The Red C12 particles Area in LD were quantified with the ImageJ ‘‘analyze particles’’ function in threshold images, with size (square μm) settings from 0.1 to 100 and circularity from 0 to 1. To determine Red C12 % area per cell, Red C12 particles in LD area was normalized with Cell area calculated with Cellpose generalist algorithm for cell segmentation.

LD (BODIPY 493/503) or Lysosome (LysoTracker™ Red DND-99) particle area was quantified with the ImageJ ‘‘analyze particles’’ function in threshold images, with size (square μm) settings from 0.1 to 100 and circularity from 0 to 1. To determine LD or Lysotracker % area per cell, LD or lysosomal particles were normalized with Cell area. Cell area was automatically generated using Cellpose generalist algorithm for cell segmentation.

### SeaHorse XF-96 metabolic flux analysis

For Seahorse metabolic flux experiments, oxygen consumption rates and extracellular acidification rates were measured using a 96-well Seahorse Bioanalyzer XF 96 according to the manufacturer’s instructions (Agilent Technologies).

#### XF Cell Mito Stress Test

Primary fibroblast were seeded (18 000 cells per well) into 96-well plates the previous day to the assay. Prior to assay, STV condition cells were switched to STV medium for 6h before analysis. Analysis was performed according to the manufacturer’s instructions. 1h prior to analysis, media was replaced with 175uL Mito assay medium and incubated for 1h at 37°C without CO2. During the experiments inhibitors were sequentially injected: Oligomycin [2uM], FCCP [2uM], Rotenone [0.5uM] and Antimycin [0.5uM]. Then OCR was automatically calculated by the Seahorse XF-24 analyzer (Seahorse Bioscience, CA, USA).

ATP production was calculated as the difference between the last rate measurement before Oligomycin injection and the minimum rate measurement after Oligomycin injection. Mitochondrial maximal respiration calculated as the difference between the maximum rate measurement after FCCP injection and the non-mitochondrial respiration. Mitochondrial spare respiratory capacity calculated as the difference between the maximal and basal respiration.

Mito assay medium: 8.7g/L MEM (61100-087, ThermoFisher scientific), 1mM pyruvate (GIBCO), 2mM glutamine and 10mM glucose, pH=7.4.4

#### The XF Palmitate Oxidation Stress Test

At 48 h before the experiment, cells were cultured on XF-96 plates at a density of 18.000 cells/well. At 24 h before metabolic flux analysis, the culture medium was replaced with 100 μL of substrate-limited medium. Before the analysis, the medium was replaced with 135 μL FAO Assay Medium and incubated at 37°C in a non-CO2 incubator for 1h. Just prior to assay cells were treated with palmitate-BSA (200 μM) or BSA (34 μM), and during the experiment inhibitors were sequentially injected: Etomoxir (40 μM), Olygomycin (2uM), FCCP (2uM), antimycin A (0.5 μM) and rotenone (0.5 μM). Then OCR was automatically calculated by the Seahorse XF-24 analyzer (Seahorse Bioscience, CA, USA).

Lipidic Oxidative respiration calculated as the fold change between Etomoxir treatment and Vehicle in the difference between the second-rate measurement after Etomoxir injection and the last rate measurement before Etomoxir injection. Lipidic ATP production was calculated as the fold change between Palmitic acid treatment and BSA treatment in the difference between the last rate measurement before Oligomycin injection and the minimum rate measurement after Oligomycin injection. Lipidic maximal respiration calculated as the fold change between Palmitic acid treatment and BSA treatment in the difference between the maximum rate measurement after FCCP injection and the non-mitochondrial respiration. Lipidic spare respiratory capacity calculated as the fold change between Palmitic acid treatment and BSA treatment in the difference between the maximal and basal respiration.

Substrate-Limited Medium: Seahorse XF DMEM (103575-100, Agilent), 0.5 mM D-Glucose (141341, ITW reagents), 1mM GlutaMAX (35050038, Gibco), 05 mM L-Carnitine (C0158, Sigma Aldrich), 1% FBS (10270-106, Gibco), pH=7.4.

FAO assay buffer: Seahorse XF DMEM ((103575-100, Agilent), 0.5 mM L-Carnitine (C0158, Sigma Aldrich), pH=7.4.

### Transmission electron microscopy TEM

Primary fibroblast were fixed in 3% v/v glutaraldehyde in 0.1 M sodium phosphate buffer (pH 7.4) for 10min at 37°C and 2h at RT. The samples were washed five times in 0.1 M sodium phosphate buffer (pH 7.4). The fixed cells were delivered to the core facility of Electron Microscopy of Principe Felipe Research Center. After this, ultra-thin sections (70nm) were done to be examined under transmission electron microscopy in the core facility of polymer characterization (UPV/EHU). Lysosomes, Autophagosomes, Fingerprint structures, lipid droplet and mitochondria area were quantified in 10–15 cells per sample cristae measurement as [41].

### Statistical methods

Statistical comparisons were carried out with the GraphPad Prism 10.0.0 software. All experiments were performed in biological replicates and plotted as mean ± SEM from three independent experiments, unless stated otherwise. Statistical significance was measured by an unpaired two-tailed t test. Significance in all figures is indicated as follows: ns (non-significant), p > 0.05, *p < 0.05, **p < 0.01, ***p < 0.001, ****p < 0.0001.

## RESULTS

### Skin fibroblasts from GRN c.709-1G>A mutation carriers show pathological hallmarks that are typically observed in the central nervous system (CNS) of FTD patients

Heterozygous GRN pathogenic variants result in approximately 50% decrease in both PGRN mRNA and protein levels [9, 22]. This feature was reproduced in fibroblasts from c.709-1G>A GRN mutation carriers, displaying reduced protein levels of GRN (p<0.05)(Fig. 1A) and mRNA (p<0,05)(Fig. 1B). Based on this finding, we investigated whether this reduction could induce FTD-like pathology in fibroblasts of patients.

**Figure 1.**
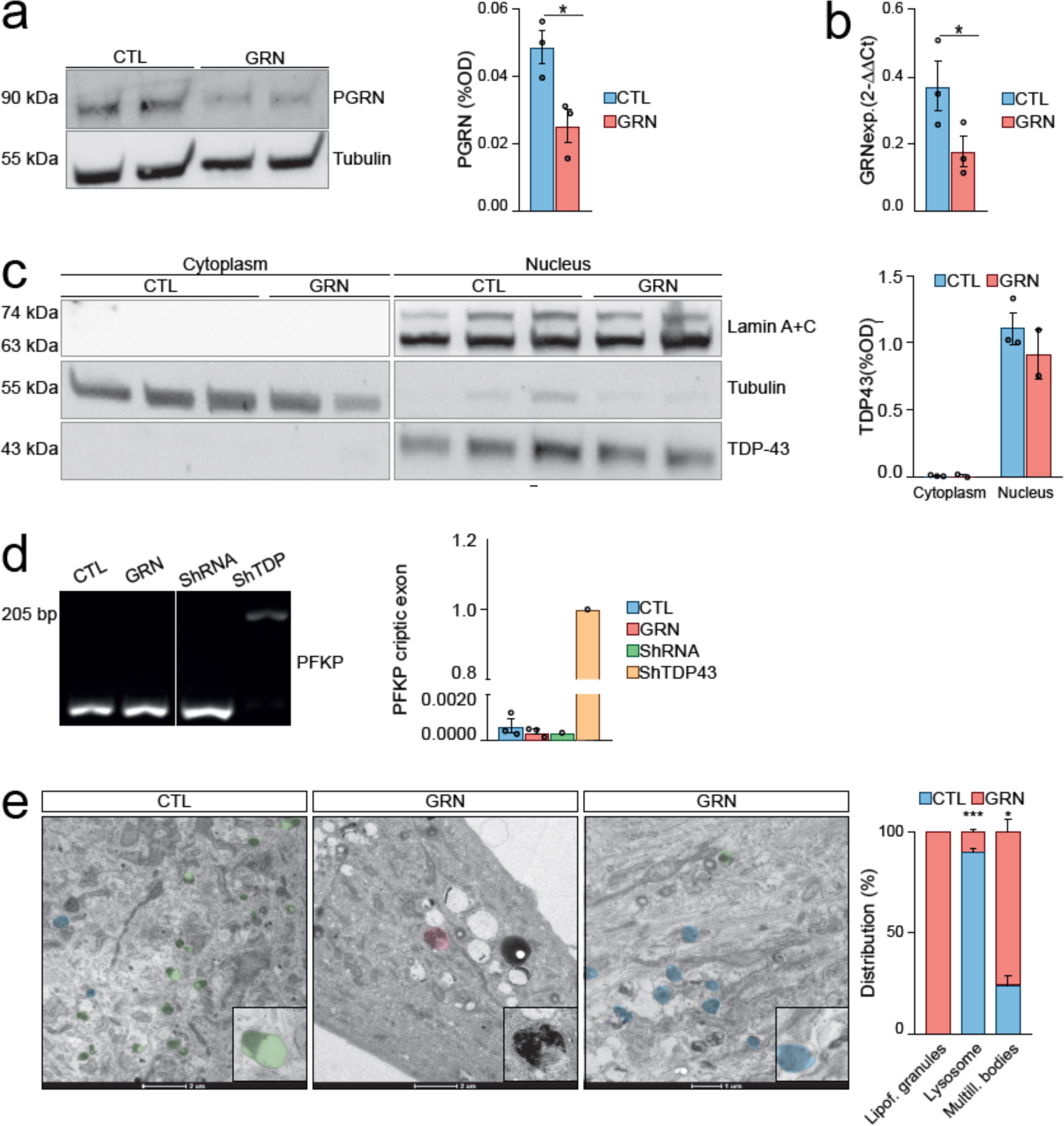
Pathological Features in Fibroblasts from Heterozygous GRNc.709-1G>A Mutation Carriers Resemble FTD Pathology. A. PGRN Protein Levels Assessed by Western Blotting in Primary Human Fibroblasts. B. PGRN mRNA Expression Analyzed by qPCR in Primary Human Fibroblasts. C. TDP-43 Protein Expression in Cytoplasmic and Nuclear Fractions of Primary Human Fibroblasts Detected via Western Blotting. D. Agarose Gel Electrophoresis of PFKP PCR Product from Primary Human Fibroblasts to assess cryptic exon retention as a surrogate marker of TDP-43 loss-of-function. E. Representative TEM Images of Primary Human Fibroblasts showing Lysosomes (green), multilamellar bodies (blue), and lipofuscin body (pink). Insets: (a) primary lysosome (electron-dense and homogeneous) and (b) secondary lysosome (heterogeneous structure). Note that the membrane-bound in both lysosomes. (c) Lipofuscin body (d) multilamellar body. Scale bar is indicated in the figure. Healthy control fibroblasts (CTL), FTD-GRN patient fibroblast (GRN), expression (Exp.). Data are presented as means ± SEMs (n = 3). Statistical significance denoted by *p < 0.05 and ***p < 0.001, determined using Student’s t test.

GRN-associated FTD is characterized by ubiquitin-positive nuclear and cytoplasmic inclusions and is composed of phosphorylated TDP-43. These aggregates are insoluble and accumulate in the FTD-affected brain regions of FTD-GRN patients. Additionally, they have been detected in cellular models where GRN has been stably silenced, and occasionally in aged *Grn*-knockout mice [34, 39, 58, 82, 85]. We performed an immunoblot on cytoplasmic and nuclear fractions of fibroblasts to investigate whether the reduction in GRN levels is sufficient to promote TDP-43 pathology. TDP-43 was found predominantly in the nuclear fraction of either healthy or FTD-GRN fibroblasts (Fig. 1C). While cytoplasmic accumulation of TDP-43 was not detected, we sought to investigate the potential loss of TDP-43 function which represents an early molecular alteration that precedes the formation of cytoplasmatic aggregates [80]. To achieve this, we conducted PCR-based examination of cryptic exons in Phosphofructokinase Platelet (*PFKP*), which are characteristic of TDP-43 loss of splicing function. However, both FTD-GRN and healthy control fibroblasts exhibited the absence of the PFKP splicing variant, which was expected at 205bp, as demonstrated in a sample of SH-SY5Y cells with lentiviral TAR DNA binding protein (*TARDBP*) knockdown (Fig. 1D). Hence, GRN haploinsufficiency does not result in any changes in TDP-43 localization or function in fibroblasts from FTD-GRN patients, at least under basal culture conditions.

In addition, GRN mutation carrying FTD patients exhibit a build-up of lipofuscin granules and lysosomal deposits [79]. These pathological features are typically associated with NCL, a condition caused by the total absence of PGRN. To explore these pathological features, both healthy control and FTD-GRN patient fibroblasts were examined by transmission electron microscopy (TEM). FTD-GRN fibroblasts showed a prominent accumulation of lysosome-related storage material, with a significantly higher proportion of multilamellar bodies compared to the healthy controls (p<0,05) (Fig. 1E, detail in Fig. S5-7). Furthermore, dense lipofuscin granules, which were not present in healthy controls, were observed in FTD-GRN fibroblasts (Fig. 1E, detail in Fig. S5-7). Additionally, the number of electron-dense lysosomes was reduced in the fibroblasts of FTD-GRN patients (p<0,05) (Fig. 1E, detail in Fig. S6-7). Next, we proceeded to quantitatively validate these presumably lysosomal defects and investigate whether they are a direct consequence of PGRN deficiency.

### Fibroblasts from GRN c.709-1G>A mutation carriers exhibit impaired lysosomal function

We conducted several analyses to confirm that the accumulation of lysosome-related storage material in fibroblasts of patients is a result of lysosomal dysfunction. Initially, we assessed the lysosomal load by measuring the mRNA and protein levels of two structural lysosomal proteins, lysosome-associated membrane glycoprotein 1 (LAMP-1) and lysosome-associated membrane glycoprotein 2 (LAMP-2), and mRNA levels of lysosomal associated membrane protein 2 (LAMP2) detecting no differences between FTD-GRN and healthy control fibroblasts (Fig. 2A-B). Additionally, there were no differences in lysosomal localization between the patient and control fibroblasts as determined by immunofluorescence (Fig. 2C). However, we observed a significant reduction in the staining of Lysotracker Red DND-99, a dye to label acidic organelles in live cells, in the patients’ fibroblasts (p<0.05). After six hours of STV, the number of active lysosomes increased as expected in healthy control fibroblasts (CTL) (CTL_NT vs CTL_STV p<0,05) but not in FTD-GRN patient fibroblasts (GRN). However, in both healthy control and patient fibroblasts, treatment with bafilomycin A1 (BafA1), an inhibitor of Vacuolar H+-ATPase, resulted in inhibition of lysosomal function (CTL_STV vs CTL_STV_BafA1 p<0,01; GRN_STV vs GRN_STV_BafA1 p<0,05) (Fig. 2D).

**Figure 2.**
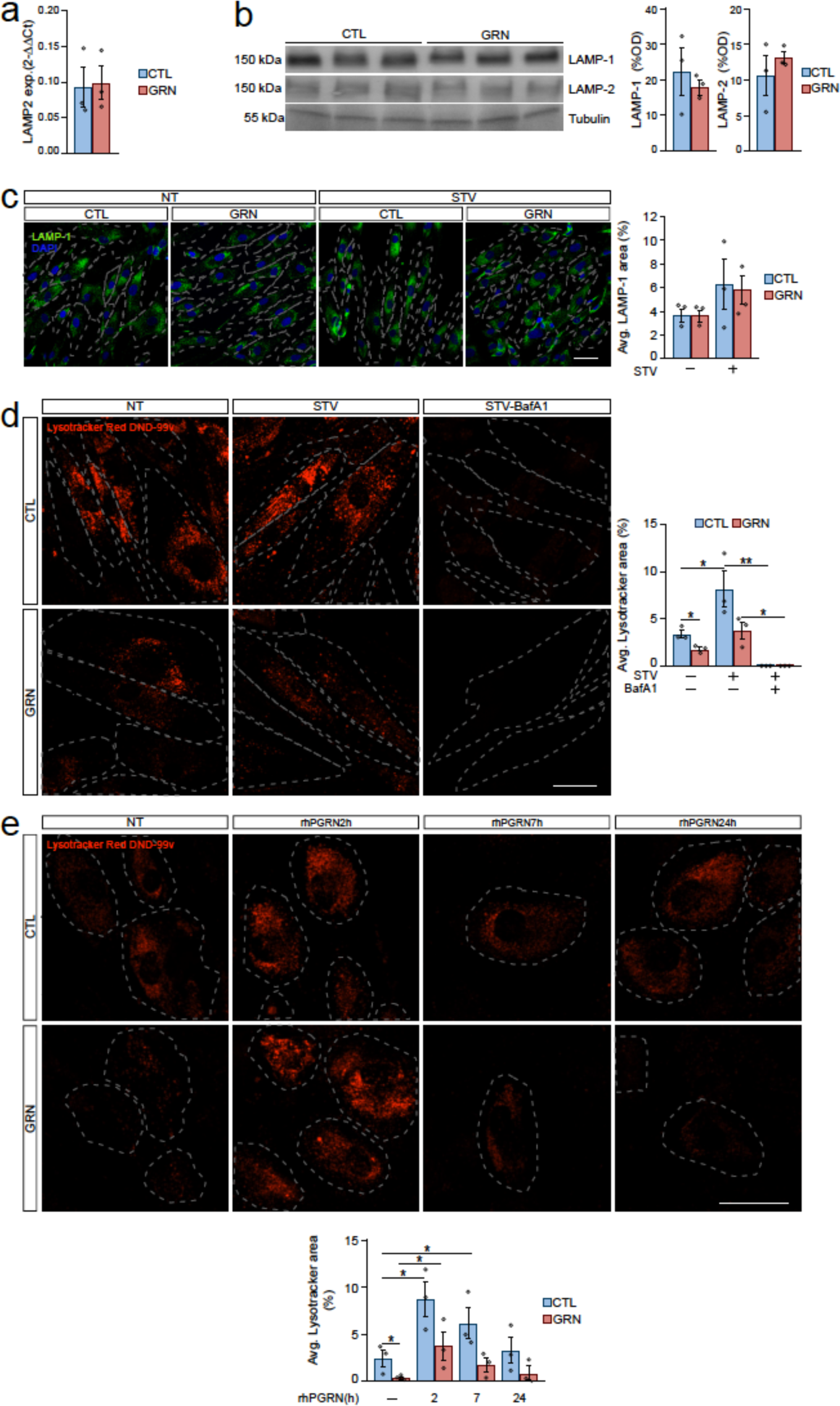
Impaired Lysosomal Function in Fibroblasts from Heterozygous GRNc.709-1G>A Mutation Carriers. A. mRNA Expression of LAMP2 Assessed by qPCR in Primary Human Fibroblasts. B. Protein Expression of LAMP-1 and LAMP-2 Analyzed by Western Blotting in Primary Human Fibroblasts. C. Representative Images of LAMP-1 (green) and DAPI (blue) staining in Primary Human Fibroblasts, cultured with or without a 6-hour pre-assay period under STV conditions. Scale bar = 50 µm. D. Representative Images and quantification of Lysotracker Red DND-99 Staining in Primary Human Fibroblasts. Primary human fibroblasts were cultured with or without a 6-hour pre-assay period under STV conditions, followed by 2 hours of incubation with BafA1 at 100 nM prior to live imaging. Lysotracker Red DND-99 staining was used to visualize lysosomal compartments. Scale bar = 40 μm. E. Representative Images and quantification of Lysotracker Red DND-99 Staining in Primary Human Fibroblasts. Primary human fibroblasts were treated with rhPGRN at 500ng/mL at different time points (2, 6, 24h). Lysotracker Red DND-99 staining was used to visualize lysosomal compartments. Scale bar = 50 µm. Healthy control fibroblasts(CTL), FTD-GRN patient fibroblast (GRN), expression (exp.), average (Avg.), non treated (NT), starving (STV), Bafilomycin A1 (BafA1), recombinant human progranulin (rhPGRN). Data are presented as means ± SEMs (n = 3). Statistical significance indicated by *p < 0.05, determined using Student’s t test.

This analysis unveiled an impaired lysosomal acidification in the fibroblasts from FTD-GRN patients, potentially explaining the observed buildup of storage material in TEM images. Since lysosomal dysfunction may be caused by insufficient PGRN, we treated the FTD-GRN fibroblasts with 500 ng/mL of recombinant human progranulin (rhPGRN) and assessed Lysotracker Red DND-99 staining at 2, 6 and 24 hours to support lysosomal acidification [73]. The peak of lysotracker staining manifested 2 hours after the treatment, observed consistently across both control and FTD-GRN fibroblasts (p<0.05) (Figure 2E). Notably, this peak in lysotracker levels reaches a magnitude comparable to that induced by a state of STV, as depicted in Figure 2D. Intriguingly, the rhPGRN treatment in FTD-GRN fibroblasts resulted in the reinstatement of functional lysosome levels comparable to those observed in untreated, healthy controls (Fig. 2E). This observation suggests a direct correlation between PGRN activity and lysosomal acidification and consequently, function in fibroblasts.

### Skin fibroblasts from FTD patients with GRN c.709-1G>A mutation display abnormalities of autophagy and mitophagy

Lysosomes play a pivotal role in the resolution of the autophagy to facilitate the enzymatic degradation of the cellular cargo. When fibroblasts are cultured under complete conditions (rich in nutrients), they do not show a significant level of basal autophagic activity. This observation is reinforced by the minimal presence of lipidated microtubule-associated protein 1A/1B-light chain 3 (LC3B-II), a reliable marker of autophagy-related structures. Consequently, these findings suggest that autophagy is not essential under basal conditions in fibroblasts (Fig. S1A). Consequently, under basal conditions it becomes challenging to detect a potential alteration in autophagy due to GRN haploinsufficiency. On the contrary, under nutrient deprivation conditions, (by serum and/or amino acid withdrawal), prominent activation of autophagosome biogenesis is detectable. Consistently, we observed an increase in the level of LC3B-II in fibroblasts after 5 hours of lysosome inhibitor chloroquine (CLQ), and no significant difference was found between samples derived from FTD-GRN patients and controls (Fig. S1B). Moreover, fibroblasts were cultured in STV medium 6h prior to assay and treated by the CLQ to measure the autophagy flux in these cells by measuring LC3B-II levels and P62 protein, also called Sequestosome 1 (SQSTM1), a ubiquitin-binding scaffold protein that binds directly to LC3B, linking ubiquitinated proteins to the autophagic machinery to enable their degradation in the lysosome. We observed a higher accumulation of LC3B-II and SQSTM1 in FTD-GRN fibroblasts when compared to healthy fibroblasts (p<0.05) (Fig. 3A). To bolster the reliability of this finding, we conducted quantitative analysis of autophagosomes through immunofluorescence of LC3B, revealing a substantial elevation in the abundance of LC3B-II positive puncta within FTD-GRN fibroblasts (p<0.05) (Fig. 3B). These findings suggest an activation of autophagosome formation in FTD-GRN patient derived fibroblasts which could be interpreted as a compensatory feedback mechanism, akin proposed in LSDs [5]. Furthermore, analysis of TEM images revealed a striking accumulation of autophagosome structures, approximately threefold higher, in FTD-GRN fibroblasts compared to healthy fibroblasts cultured under nutrient-rich conditions (p<0.05) (Fig. 3C, detail in Fig. S8,9). This observation could be convincingly attributed to the combined effects of lysosomal not acid pH, which produces a functional impairment, and the compensatory activation of autophagosome biogenesis.

**Figure 3.**
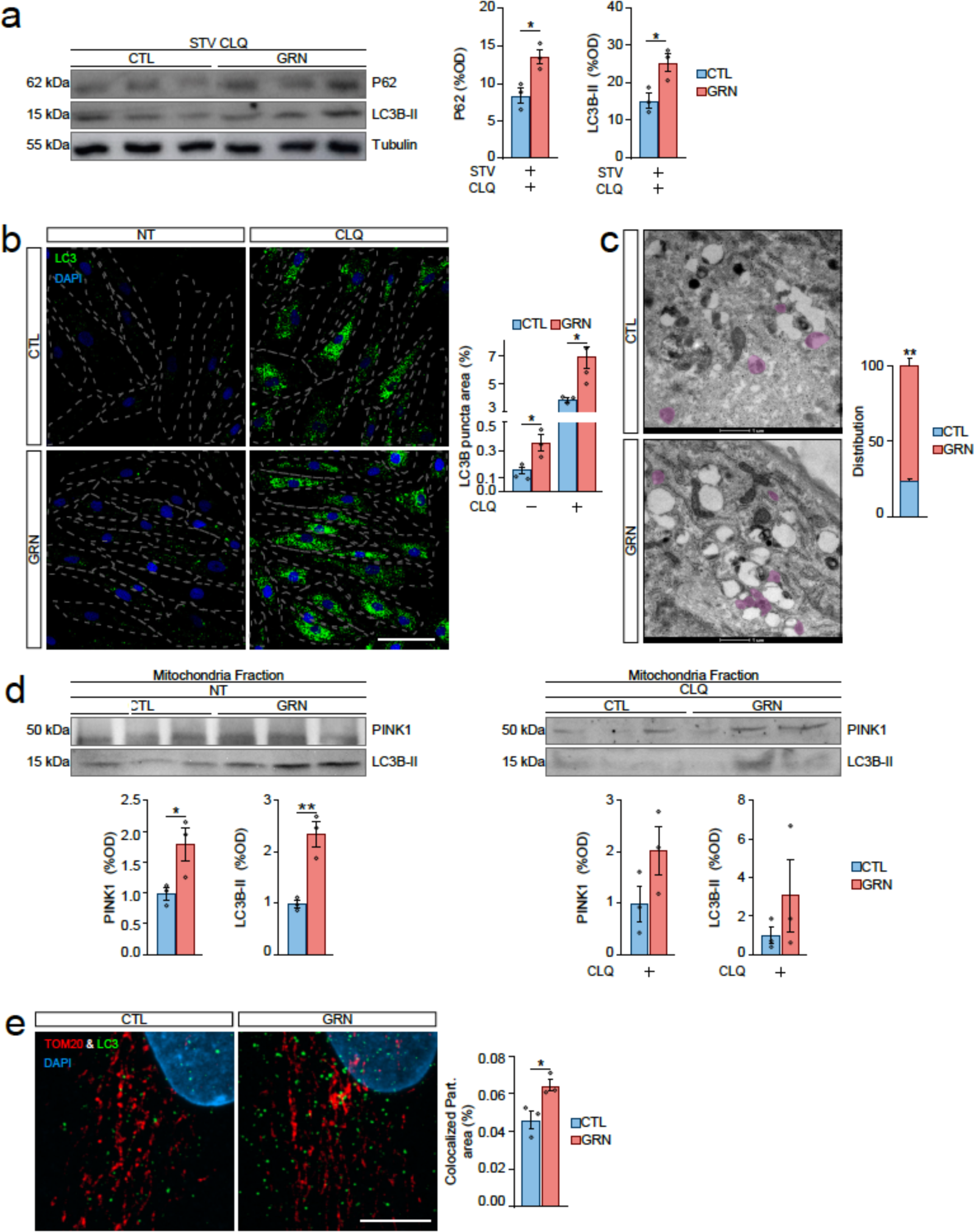
Dysregulated Autophagy and Mitophagy in Fibroblasts from GRNc.709-1G>A Mutation Carriers. A. Protein Levels of LC3B-II and P62 Assessed by Western Blotting in Primary Human Fibroblasts. Primary human fibroblasts were cultured with or without a 6-hour STV pre-assay period and treated with CLQ at 30 µM for 5 hours. Western blotting was performed to assess the protein levels of LC3B-II and P62, and quantification are shown as bar plot graphs. B. Representative Images of LC3B Puncta (green) and DAPI (blue) Staining in Primary Human Fibroblasts. Primary human fibroblasts were treated with or without a 5-hour pre-assay period with CLQ at 30 µM. LC3B puncta (green) staining and DAPI (blue) staining were used to visualize autophagosomes and nuclei, respectively. Scale bar = 100 µm. C. Representative TEM images of primary human fibroblasts, showing autophagosomes (purple). Insets: Autophagosome. Scale bar is indicated in the figure. D. Protein Levels of LC3B-II and PINK1 Analyzed by Western Blotting in Cytoplasmic and Mitochondrial Fractions of Primary Human Fibroblasts. Primary human fibroblasts were treated with or without a 5-hour pre-assay period with CLQ at 30 µM. Western blotting was performed to analyze the protein levels of LC3B-II and PINK1 in both cytoplasmic and mitochondrial fractions. E. Representative Images of LC3B (green) and TOM20 (red) Staining in Primary Human Fibroblasts. Bar plot shows the quantification of the relative area of LC3B colocalized with TOM20. Scale bar = 10 µm. Healthy control fibroblasts(CTL), FTD-GRN patient fibroblast (GRN). Starving (STV), Chloroquine (CLQ), non treated (NT). Data are presented as means ± SEMs (n = 3). Statistical significance represented by *p < 0.05 and **p < 0.01, determined using Student’s t test.

Autophagy is triggered not only in response to nutrient deprivation but also in reaction to the buildup of impaired cellular organelles like mitochondria. Hence, we investigated the potential impact of GRN haploinsufficiency on mitophagy by quantifying the levels of LC3B-II and PTEN-induced kinase I (PINK1) within the mitochondrial fraction. The elevation of PINK1 levels is indicative of compromised mitochondrial health and could guide the initiation of mitophagy. Therefore, we obtained a mitochondrial-enriched fraction which was validated by mitochondrial import receptor subunit TOM20 homolog (TOM20) mitochondrial marker presence, together with Glyceraldehyde-3-phosphate dehydrogenase (GAPDH) cytosolic marker absence immunoblotting (Fig. S2). Importantly, we detected higher levels of PINK1 in FTD-GRN patient fibroblasts, as depicted in Figure 3D (p<0.05). Alternatively, CLQ did not induce an increase in PINK1 levels, probably because CLQ halts lysosomal degradation while not influencing the initial pool of PINK1 tagged mitochondria in fibroblasts. Therefore, it can be inferred that mitophagy in fibroblasts is not a frequent process. Nonetheless, this observation hints at a slight enhancement in the autophagic induction of mitochondria in cells from FDT-GRN patients (Fig. 3D). Additionally, we noticed a significant colocalization of mitochondria (TOM20 staining) and autophagosome cargo (LC3B staining) in FTD-GRN fibroblasts (p<0,05) (Fig. 3E). Altogether, these findings suggest that mitochondria are not able to evade the enhanced autophagy flux induced by GRN haploinsufficiency. We anticipated that this phenomenon could have an impact on mitochondria, and therefore we sought to investigate the consequences arising from GRN haploinsufficiency on mitochondrial functionality.

### GRN haploinsufficiency causes mitochondrial abnormalities in GRN c.709-1G>A mutation carrier skin fibroblasts

The quantification of the mitochondrial marker TOM20, utilizing both immunoblotting and immunofluorescence methodologies, revealed no discernible impact on either the mitochondrial surface area or the integrity of the mitochondrial network. This observation is substantiated by the comparable lengths of mitochondrial branches, as shows Fig. 4A. The ultrastructure assessment using TEM unveiled a pronounced escalation in the prevalence of swollen mitochondria (p<0.05) and a concomitant reduction in the abundance of mitochondrial cristae within FTD-GRN fibroblasts when compared to the control group (p<0.005) (Fig. 4B, detail in Fig. S10,11). To probe the functional power of these mitochondria, we assessed mitochondrial respiration utilizing SeaHorse technology. Our findings elucidate that the basal ATP production might be diminished in FTD-GRN fibroblasts, although this analysis did not achieve statistical significance due to notable variability of control fibroblasts (Fig. 4C). However, a substantial impairment in maximal respiration capacity and spare respiratory capacity was distinctly evident in FTD-GRN fibroblasts (p<0.05), attributed to a concurrent reduction in spare respiration (p< 0.05) (Fig. 4C). Mitochondrial respiration was additionally evaluated under conditions of cellular STV (Fig. S. 3A). To elucidate the potential modulation of mitochondrial respiratory capacity by PGRN, we treated fibroblasts with 500 ng/mL of rhPGRN for varying durations. Our findings revealed that rhPGRN treatment displayed the capacity to enhance mitochondrial ATP production and elevate maximal respiratory capacity and spare respiratory capacity particularly in FTD-GRN fibroblasts. Interestingly, this effect did not manifest at the 2-hour time point following treatment initiation, a period during which rhPGRN did exhibit enhancements in lysosomal function (Fig. 3A). Instead, discernible improvements were evident in later time intervals, specifically within the range of 6 to 12 hours (Fig. 4D). In conjunction with the observed time-dependent effects, our investigation also unveiled dose-dependent responses. Notably, the application of rhPGRN at a concentration of 1000 ng/mL yielded a more pronounced enhancement in ATP production capacity already at 6 hours following treatment initiation (Fig. 4E). However, it is noteworthy to mention that this augmentation did not correspondingly impact the maximal respiratory capacity or spare respiratory capacity (Fig. 4E). These findings present compelling evidence indicating pronounced abnormalities in the mitochondrial network within FTD-GRN fibroblasts. This aberration is concomitantly associated with a compromised respiratory capacity, which, albeit exhibiting a modest effect size, is mitigated through intervention with rhPGRN treatment.

**Figure 4.**
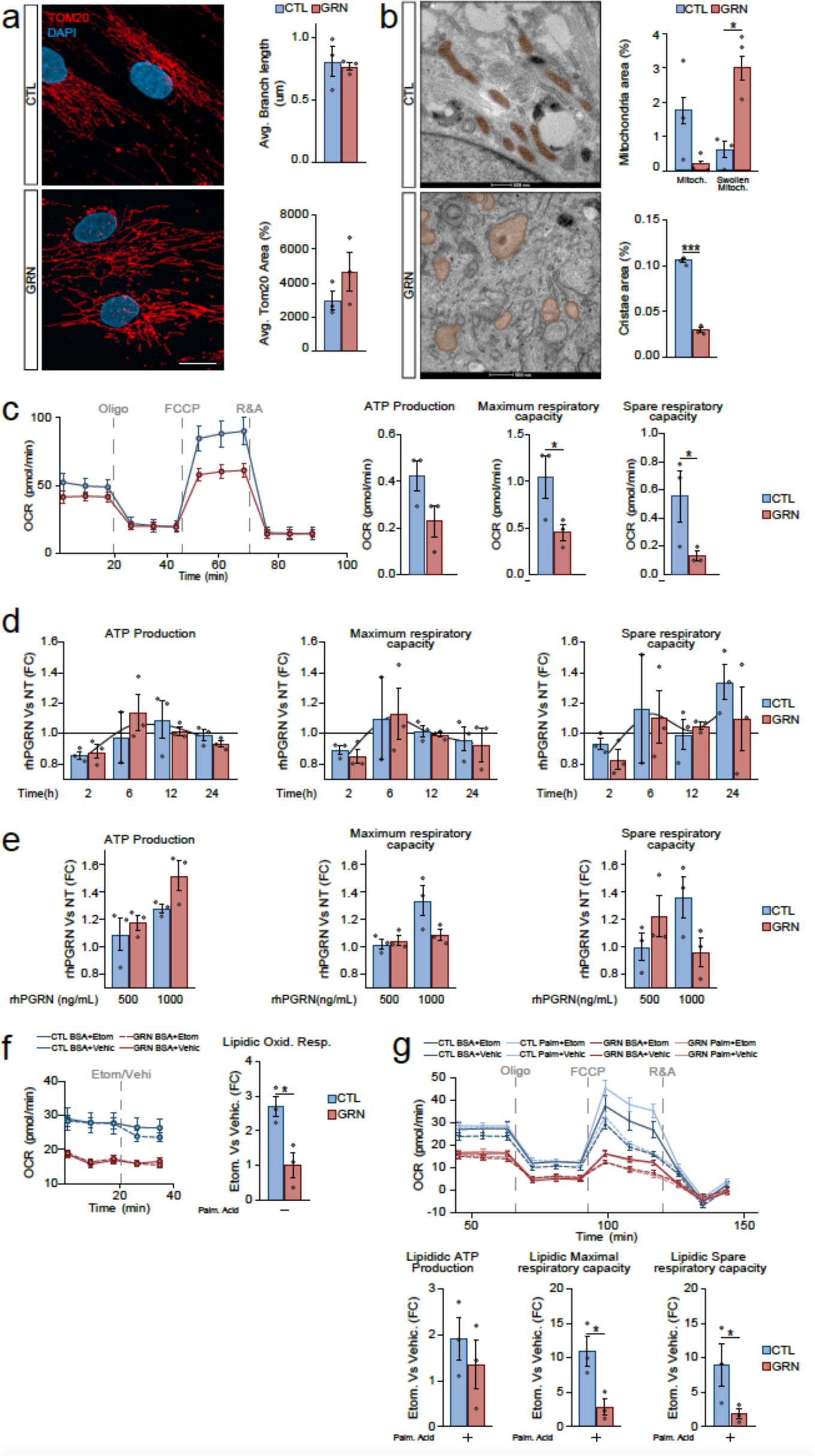
Altered Mitochondrial Function in Fibroblasts from Heterozygous GRNc.709-1G>A Mutation Carriers. A. Representative Images of TOM20 (red) and DAPI (blue) Staining in Primary Human Fibroblasts. Scale bar = 20 µm. B. Representative TEM Images of Primary Human Fibroblasts showing mitochondria ultrastructure (orange). Insets: (a) Functional mitochondria, (b) swollen mitochondria. Scale bar is indicated in the figure. C. Seahorse Assay depicting Mitochondrial Oxygen Consumption in Primary Human Fibroblasts. D. Seahorse Assay measuring Mitochondrial Oxygen Consumption in Primary Human Fibroblasts, cultured with or without the specified pre-assay period, and treated with rhPGRN at 500 ng/mL at different time points (2, 6, 12 and 24h). E. Seahorse Assay analyzing Mitochondrial Oxygen Consumption in Primary Human Fibroblasts, cultured with or without a 6-hour pre-assay period, and exposed to rhPGRN at 500ng/mL or 1000ng/mL. F. Seahorse Assay Assessing Mitochondrial Free Fatty Acid Oxygen Consumption in Primary Human Fibroblasts. The left panel illustrates the time-course evolution of the basal oxygen consumption rate (OCR). The right panel presents the quantification of lipidic oxidative capacity, represented as the change in OCR induced by etomoxir treatment. G. Seahorse Assay Measuring Mitochondrial Free Fatty Acid Oxygen Consumption in Primary Human Fibroblasts. The left panel illustrates the time-course evolution of the oxygen consumption rate (OCR) during the assay. The right panel presents the quantification of lipidic ATP production, lipidic maximal respiratory capacity and lipidic spare respiratory capacity, represented as the change in those parameters induced by etomoxir treatment. Healthy control fibroblasts (CTL), FTD-GRN patient fibroblast (GRN), average (Avg.), Mitochondria (Mit.), oligomycin (Oligo), rotenone and antimycin (R&A), recombinant human progranulin (rhPGRN), non treated (NT), fold change (FC), etomoxir or vehicle (Etom/Vehi), palmitic acid (Palm. Acid) Data are presented as means ± SEMs (n = 3). Statistical significance represented by *p < 0.05 and **p < 0.01, determined using Student’s t test

Our aim was to investigate whether the observed mitochondrial defects in FTD-GRN fibroblasts specifically impact any energy source. Interestingly, subjecting both FTD-GRN and control fibroblasts to a 6-hour amino acid deprivation prior to SeaHorse analysis resulted in no significant alteration in mitochondrial respiratory capacity (Fig. S3A). This suggests that fibroblast mitochondria may not heavily rely on amino acids as a primary energy source or that GRN haploinsufficiency does not exert substantial net effects on amino acid-driven respiration. Conversely, when the beta-oxidation of fatty acids (FAO) was hindered through etomoxir treatment, an irreversible inhibitor of Carnitine O-palmitoyltransferase 1 (CPT1) that impedes the transportation of FAs into the mitochondria, lipidic oxygen consumption, maximal lipidic respiration capacity and lipidic spare respiration capacity, ATP production and maximal respiration capacity declined in control fibroblasts (p<0.05) but remained unaffected in FTD-GRN fibroblasts (Fig. 4F,G). This observation suggests a plausible impairment in mitochondrial FAO under conditions of GRN haploinsufficiency, consequently leading to an elevated likelihood of lipid accumulation.

### GRN insufficiency causes impairments of lipid metabolism in skin fibroblasts patients with GRN c.709-1G>A mutation

Accordingly, our focus shifted towards intracellular organelles termed LDs, acting as reservoirs for stored lipids destined for future utilization. The analysis of LDs by TEM images unveiled a notable rise of these structures in FTD-GRN fibroblasts (p<0.05), often situated proximal to or in direct contact with the ER and characterized by an electron-lucent core encompassed by an electron-dense periphery (Fig. 5A, detail in Fig. S12,13). This pattern is consistent in FTD-GRN fibroblasts where the incorporation of newly synthesized TAGs which are more electron-dense, into pre-existing LDs enriched with cholesterol esters which are electron-lucent [20], resulting in a liquid-to-liquid phase separation (LLPS) of these two lipid components. This discovery prompted us to delve into the way the regulation of LD dynamics is influenced in FTD-GRN cells, and to elucidate whether treatment with rhPGRN can exert a modulatory influence on this process. To monitor the recent incorporation of esterified FAs into LDs, we implemented a pulse-chase assay utilizing BODIPY 558/568 C12 (Red-C12) (Fig. 5B). This compound is acknowledged for its conversion into neutral lipids, making it a valuable tool for studying lipid trafficking [67]. The dynamics of LDs are intricately regulated, especially in conditions of nutrient deprivation. During such periods, FAs can be mobilized from membrane-bound organelles through autophagy and lysosomal lipolysis, thereby augmenting the LD pool. Conversely, FAs can be released from LDs themselves by lysosomal (lipophagy) and cytoplasmic lipolysis, facilitating their transfer into mitochondria [27, 59, 67]. Consequently, we conducted a comparative analysis between cells cultured under nutrient-rich conditions and to nutrient deprivation. As anticipated, our observations revealed an elevation in LD formation under nutrient-deprived conditions, a phenomenon that exhibited similarity between both control and FTD-GRN fibroblasts. This was evident through both increased BODIPY 493/503 (BD) staining of LDs and, importantly, the vast majority of Red-C12 being localized within LDs (p<0.05) (Fig. 5C). However, when nutrients were not limited FTD-GRN fibroblasts demonstrated elevated levels of BD staining in comparison to healthy controls (p<0.05), as indicated in Fig. 5C. This observation is in accordance with our TEM imaging findings (Fig. 5A) and is further supported by a larger portion of Red-C12 being localized within LDs (p<0.05) (Fig. 5C). The observed reduction in differences under nutrient deprivation might be attributed to either a saturation point being reached in the cells’ capability to store LDs, or a decreased biogenesis of LDs through autophagolysosome-driven processes in FTD-GRN fibroblasts.

**Figure 5.**
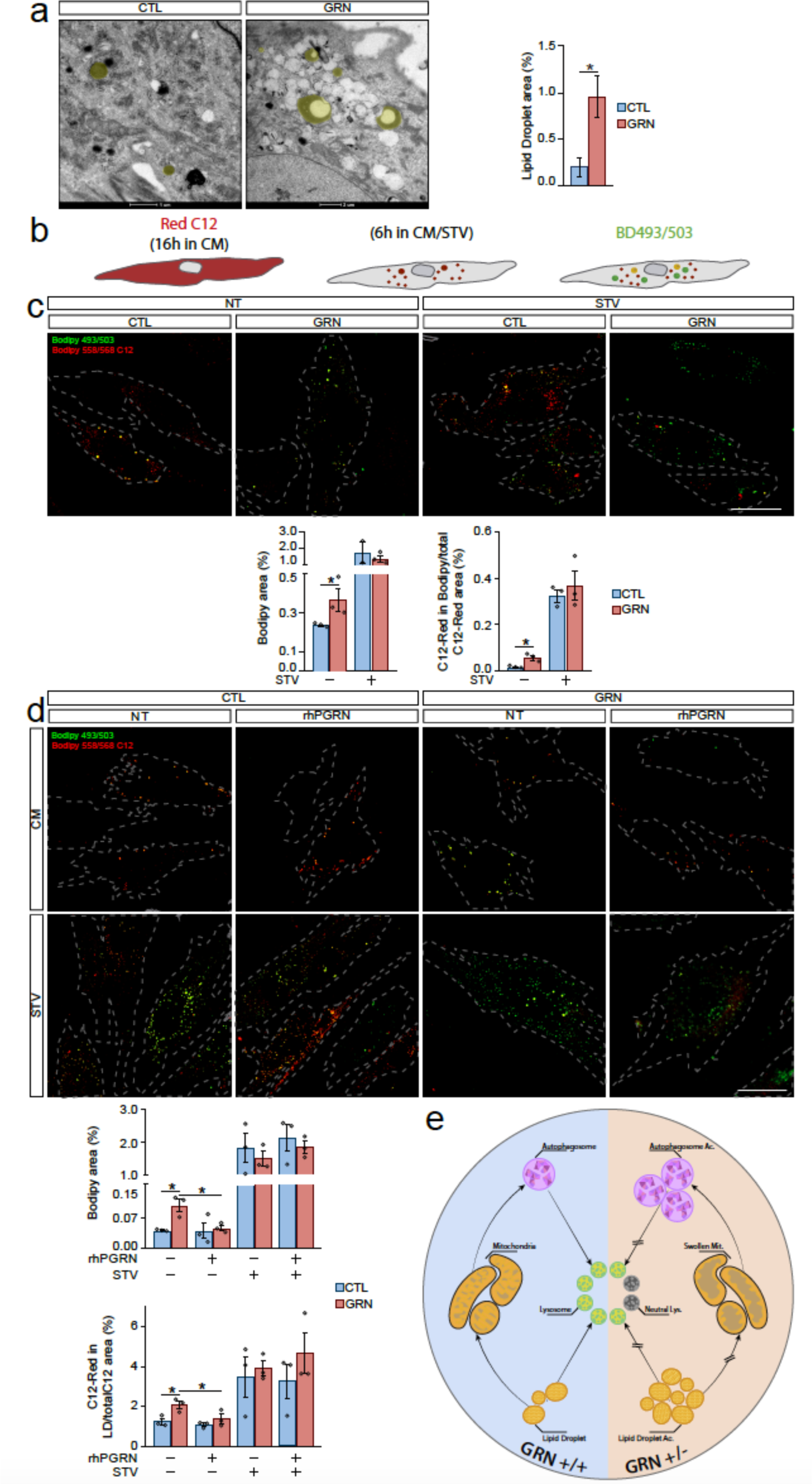
Disrupted Lipid Metabolism in Fibroblasts from Heterozygous GRNc.709-1G>A Mutation Carriers. A. Representative TEM Images of Primary Human Fibroblasts, showing LDs (yellow). Inset: High magnification of a LD with a LLPS of lipid components: electron-lucent core and electron-dense capsule. Note the direct interaction of the membraneless LD surface with a network of vimentin intermediate filaments. Scale bar is indicated in the figure. B. Schematic Depiction of the Pulse and Chase Assay. C. Representative Images of BD (green) and Red C12 (red) Staining in Primary Human Fibroblasts cultured under normal or STV conditions (6h). Bar plots show quantifications of area covered by BD and the proportion of C12 staining within BD-positive LDs. Scale bar = 50 µm. D. Representative images of BD (green) Staining in Primary Human Fibroblasts cultured under normal or STV conditions, and treated with rhPGRN at 500ng/mL for 2h. Bar plots show quantification of area covered by BD and the proportion of C12 staining within BD-positive LDs.Scale bar = 50 µm E. This graphical abstract provides a simplified cellular landscape representing the pathological mechanism connecting GRN insufficiency with lipid dysregulation. The decrease in GRN levels leads to the accumulation of ineffective lysosomes, which directly impacts mitochondrial dynamics, potentially compromising their renewal process. This mitochondrial dysfunction subsequently disrupts fatty acid processing, culminating in lipid accumulation and, most likely, contributing to lipotoxicity. Healthy control fibroblasts(CTL), FTD-GRN patient fibroblast (GRN). Starving (STV), recombinant human progranulin (rhPGRN), accumulation (Ac.), lysosome (Lys.), mitochondria (Mit.). Data are presented as means ± SEMs (n = 3). Statistical significance denoted by *p < 0.05, determined using Student’s t test.

To ascertain whether the observed alterations in LD dynamics within FTD-GRN fibroblasts are directly attributed to GRN insufficiency, we treated both control and FTD-GRN fibroblasts with rhPGRN for 2h, at 500 ng/mL. As depicted in Fig. 5D, the 2-hour rhPGRN treatment exhibited a significant reduction in LD accumulation within FTD-GRN fibroblasts, effectively restoring up to control fibroblasts levels (p<0.05). It is noteworthy that this effect was specific to FTD-GRN fibroblasts, as rhPGRN treatment did not induce any alteration in LD levels within healthy control fibroblasts (Fig. 5D). The rhPGRN treatment led to a reduction in the proportion of Red-C12 within LDs (p<0.05), bringing them to levels comparable to controls under nutrient-rich conditions (Fig. 5D). However, this treatment did not induce any changes under nutrient-deprived conditions (Fig. 5D). In any case, under nutrient-rich conditions, LD dynamics appear to be directly influenced by GRN haploinsufficiency in FTD-GRN cells. As posed before, the mechanisms governing these dynamics may involve both autophagic and mitochondrial processes, both of which have been demonstrated to be impaired by PGRN deficiency. Indeed, concerning mitochondria, we have observed a similar accumulation of Red-C12 within mitochondria of FTD-GRN fibroblasts, as indicated by Red-C12 staining in Mitotracker-positive structures (Fig. S4), suggesting that the increased levels of LD could, in part, stem from a potential impairment in FAO.

## DISCUSSION

Our study investigated the cellular mechanisms underlying FTD in GRN c.709-1G>A mutation carriers. We found that GRN insufficiency leads to a complex interplay of cellular changes in FTD-GRN fibroblasts, which provides new insights into the disease’s underlying mechanisms.

The comparison of fibroblasts from FTD-GRN patients and healthy controls revealed the presence of several key pathological hallmarks, akin to those observed in the central nervous system of FTD patients. First, we confirmed the reduction of GRN protein and mRNA levels in FTD-GRN fibroblasts, consistent with the haploinsufficiency characteristic of GRN-associated FTD [9, 22]. Secondly, discernible impairment in lysosomal acidification and perturbed regulation of autophagic flux were evident, potentially underpinning the observed accumulation of storage material, a phenomenon well-substantiated within the cerebral cortex of FTD-GRN patients [73]. Of notable significance, administration of rhPGRN exhibited efficient restoration of lysosomal acidification, thereby elucidating a direct nexus between PGRN activity and fibroblast lysosomal activity. These findings guarantee the applicability of fibroblasts as a legitimate model for investigating the etiopathological basis of FTD. Accordingly, previous works have documented the presence of accumulated storage material alongside lysosomal aberrations in patient-derived cellular cultures derived from extra-neuronal tissues, including fibroblasts [79] as investigated in our present study, and lymphoblasts [3].

We also investigated the impact of GRN insufficiency on the development of TDP-43 pathology in fibroblasts, a hallmark feature of FTD. Notably, GRN haploinsufficiency displayed no noticeable impact on the localization or splicing function of TDP-43 under basal culture conditions, implying that aberrant modulation of TDP-43 activity may not be a prerequisite for instigating lysosomal dysfunction and the accumulation of storage material. It is important to underscore that this observation does not imply that TDP-43 pathology is an epiphenomenon devoid of repercussions on neurons of FTD-GRN patients. In fact, it is widely acknowledged that alterations in TDP-43 function linked to GRN deficiency might be a pivotal contributor to the selective vulnerability of neurons [57]. Therefore, the beneficial impact of PGRN treatment, which reverses axonal damage in neurons afflicted by *TARDBP* mutations [13, 40], provides compelling support for the role of TDP-43 in this context. Though, both GRN haploinsufficiency and TDP-43 pathology synergistically could converge to trigger age-associated FTD neurodegeneration. However, complete GRN deficiency in the absence of TDP-43 pathology can independently elicit neuronal damage manifested as NCL, albeit at an exceptionally premature stage [1, 30, 74]. Hence, the absence of TDP-43 pathology recapitulation in fibroblasts from FTD-GRN patients does not diminish the significance of this model for research purposes. On the contrary, this model is effective in addressing deficiencies that are more closely associated with GRN physiology, such as lysosomal function and its derived metabolic implications. It also addresses related aspects of the disease, such as mitochondrial anomalies and their consequences, which aligns with the aims of the current study, discussed below.

A key finding of our study was that GRN haploinsufficient fibroblasts showed both mitochondrial and lysosomal structural and functional abnormalities. The mitochondria had damaged cristae, which are the inner membrane folds that are important for energy production, and consequently impaired energy production through oxidative phosphorylation (OXPHOS). A convincing link between the mitochondrial abnormalities and lysosomal dysfunction related to GRN deficiency is the deficit in mitochondrial recycling and surveillance mechanisms. This can lead to the accumulation of damaged mitochondria, which can impair overall mitochondrial function. Our study of mitophagy supports this hypothesis. In fact, mitochondrial dysfunction is becoming a major contributor to the development of LSDs, and impaired mitophagy is a common mechanism that is shared by many of these disorders [43, 71, 75]. This suggests that the imbalance between lysosomal and mitochondrial functions caused by GRN insufficiency could represent a shared mechanism contributing to the onset of neurodegenerative conditions, as has already been proposed [24, 61]. For instance, mutations that have been linked to several types of Charcot-Marie-Tooth syndrome, Parkinson’s Disease, and LSDs, have been shown to disrupt the interactions between mitochondria and lysosomes [21].

Since the disruption of the balance between lysosomes and mitochondria is likely to trigger a chain reaction of cellular events, we investigated whether this could ultimately lead to the development of a wider range of pathological changes in the context of GRN deficiency, detecting a LD accumulation in fibroblasts from FTD-GRN patients. Thus, we focused on how lipid homeostasis could be affected by this disruption. More specifically, our observations revealed a retention of FAs within LDs in fibroblasts derived from FTD-GRN patients, a phenomenon rectified upon exogenous treatment with rhPGRN. It is common for patients with neurodegenerative disorders, including FTD, to have abnormal levels of LDs in their nerve cells [2, 49, 52]. These LDs are not harmful in themselves, but they are a sign that something is wrong with the way lipids are being processed and distributed in the cells [60]. A key part of this process indeed involves the interaction between mitochondria and lysosomes [15]. Lysosomes break down lipids from cellular membranes and other sources, and provide the FAs that mitochondria need for energy [67]. If the mitochondria or lysosomes are not working properly, this process can break down, leading to the accumulation of FAs inside and outside the mitochondria. In fact, the impairment of FA b-oxidation in disrupted mitochondria might promote the cytosolic accumulation of FAs within LDs. We have demonstrated in fibroblasts from FTD-GRN patients, where decreased mitochondrial FA oxidation (FAO) was observed. This can further damage the mitochondria and make it less efficient [33, 56]. Additionally, this disruption can trigger the buildup of lipid-laden membranes within the lysosomes, which can lead to the malfunctioning of lysosomes and the subsequent accumulation of lipid storage material, as it occurs in lysosomal lipid storage diseases [37, 69]. This phenomenon has been documented by us and other researchers in fibroblasts and patients [79]. Our results suggest that cellular lipid imbalance emerges as a significant outcome in cases of lysosome-mitochondria crosstalk disruption, potentially giving rise to cytotoxic effects that may play a crucial role in the pathogenesis of FTD-GRN.

In conclusion, our study illuminates the multifaceted cellular mechanisms underlying FTD-GRN pathology. The observed lysosomal dysfunction, mitochondrial abnormalities, altered autophagy/mitophagy, and lipid metabolism impairments collectively contribute to the disease’s complexity. Moreover, based on GRN haploinsufficiency-induced lysosomal dysfunction, this study distills a pathological metabolic interplay between lysosome and mitochondria through lipid metabolism in the pathogenesis of FTD-GRN. These findings highlight potential targets for therapeutic interventions and underscore the importance of further research to fully elucidate the pathogenesis of FTD-GRN and develop effective treatments.

## Conclusion

Haploinsufficiency of GRN gene, in addition to the decreased lysosomal function, induces alterations in the cell metabolism, affecting both mitochondrial and lipid homeostasis. These metabolic alterations, arising as secondary manifestations of GRN-related lysosomal dysfunction, offer valuable insights into the fundamental pathological processes driving disease progression. These findings unveil novel potential therapeutic targets, thereby contributing toward the development of future efficacious treatments for patients with Frontotemporal dementia due to GRN mutation.

## DATA AVAILABILITY

Data sharing is not applicable to this article as no datasets were generated or analysed during the current study.

All data generated or analyzed during this study could be provided under request.

## COMPETING INTERESTS

The authors declare that they have no competing interests

## FUNDING

This work was funded by Instituto de Salud Carlos III (ISCIII) and co-funded by the European Union - European Regional Development Fund (P18/01066, PI19/01637, PI21/00153); by CIBERNED (CIBER of Neurodegenerative Diseases, CB06/05/1126, Group 609); by Diputación Foral de Gipuzkoa (projects 2021-CIEN-000020-01); by EiTB Maratoia (BIO17/ND/023/BD); by Fundació La Marató (202006-32), by IKUR strategy from Government of Basque Country (NEURODEGENPROT project) and Eusko Jaurlaritzako Osasun saila (2017222027, 2018111042, 2019222020, 2021333050,). J.O., J.L.Z-E, M.Z., and M.G-A. received support from the Department of Education of the Basque Country through grants PRE2018_1_0095, PRE2022_1_0315, PRE2015_1_0023, and PRE2020_1_0122, respectively. F.J.G.-B. was funded by the Roche Stop Fuga de Cerebros program (BIO19/ROCHE/017/BD), G.G. benefited from the Juan de la Cierva-Incorporación fellowship (Ministerio de Ciencia e Innovación, IJC2019-039965-I), and I.J., F.J.G-B., and G.G. received support from the IKERBASQUE Research Foundation.

## AUTHORS’ CONTRIBUTION

GG and FGB designed the study. JO, JZE and MZufiria, performed experiments with FTD-GRN patient derived fibroblasts of GRN expression and lysosomal function. JO, MZ and JZE performed SeaHorse, mitochondria and mitophagy experiments with help from IJH for interpretation. JO and MGA performed Lipid droplet and dynamics experiments. Transmission electron microscopy experiments were designed by FGB and GG, and performed by JO with additional support for data interpretation from ML and JR. ALM, FM and MZulaica clinically identified and characterised patients and collected samples. All data were analyzed by JO, interpreted by JO, FGB, GG and ML. FGB and GG supervised and coordinated all work. JO, GG, FGB, IJH, ALM and FM contributed to the preparation of the manuscript.

## ACKNOWLEDGEMENTS

The authors would like to specially dedicate this work to all patients with FTD-GRN and their families who participated in the study. The authors thank for technical and human support provided by SGIker (UPV/EHU/ ERDF, EU) in special to Ana Martínez from the core facility of polymer characterization (UPV/EHU) and to Mario Soriano of the core facility of Electron Microscopy of Principe Felipe Research Center.

## Notes

### Competing Interest Statement

The authors have declared no competing interest.

