## Supplemental Material and Methods for "Lipid Dysregulation Unveil the Intricate Interplay of Lysosomal and Mitochondrial Changes in Frontotemporal Dementia with GRN Haploinsufficiency"

To quantify the fluorescence intensity of LC3 puncta and Lamp2, the mean fluorescence intensity in LC3 puncta or Lamp2 channel was calculated across the entire image. To determine the % of Optical Density, the mean fluorescence intensity was normalized whit the image area.

To quantify the fluorescence intensity of LC3 in Mitochondria, we made a mask using the LC3 puncta channel, and then overlap with the Tom20 channel mask. The LC3 particles Area in Mitochondria were quantified with the ImageJ ''analyze particles'' function in thresholded images, with size (square µm) settings from 0.1 to 100 and circularity from 0 to 1. To determine LC3 % area per cell, LC3 paritcles in Mitochondria area was normalized with Cell area calculated with Cellpose generalist algorithm for cell segmentation.

Measures of mitochondrial area was generated using the 'analyse particles' function in Fiji (NIH) with a minimum area of 0.25 mm. Measures of mitochondrial length, junctions (voxels with three or more neighbors) and branches (slab segments connecting end points to either junctions or other endpoints) were determined using the 'skeletonize' and 'analyse skeleton' plugins in Fiji (NIH). Analyses were carried out on whole cells.

To quantify the fluorescence intensity of Cytoplasmic TDP43, we made a mask using the TDP43 channel, and then remove the signal from the DAPI mask. To determine TDP43 % Optical Density per cell, TDP43 mean fluosercence intensity in the cytoplasm was normalized with Cell area calculated with Cellpose generalist algorithm for cell segmentation.

**Live Cell Imaging**

For the lysosomal functionality assay fibroblast cells were incubated with CM containing 100 nM LysoTracker™ Red DND-99 (Thermo Fisher) for 5h. Cells were then washed three times with CM and immediately images were taken.

For the FA Pulse and Chase Assay fibroblast cells were incubated with CM containing 1 mM BODIPY 558/568 C12 (Red C12, Life Technologies) for 16 hr. Cells were then washed three times with CM, incubated for 1 hr in order to allow the fluorescent lipids to incorporate into LDs, and then chased for the time indicated in CM or Starving medium in the absence or presence of rhPGRN. LDs were labeled with 200ng/mL BODIPY 493/503 (Life Technologies) immediately prior to imaging and was present during imaging.

For FA tracing into mitochondria, Fibroblast cells were incubated with CM containing 1 mM BODIPY 558/568 C12 (Red C12, Life Technologies) for 16 hr. Cells were then washed three times with CM, incubated for 1 hr in order to allow the fluorescent lipids to incorporate, and then chased. Mitochondria were labeled with 100nM MitoTracker™ Green FM (M7514, InvitrogenTM) for 30min. Cells were then washed once with CM and immediately images were taken.

All images were acquired on a spinning-disk confocal (UltraVIEW VoX; Perkin Elmer) on a Nikon Eclipse Ti microscope using an Apochromat _ 100 1.49 numerical aperture oil-immersion objective (Nikon) in a temperature-controlled chamber (37 ºC).

**Image Processing, Analysis, and Statistics**

Images were analyzed using ZEN Blue Imaging Software (Zeiss) and ImageJ (NIH). Image brightness and contrast were adjusted in Adobe Photoshop CS. To quantify the fluorescence intensity of Red C12 in LDs, we made a mask using the BODIPY 493/503 channel, and then overlap with the Red C12 channel mask. The Red C12 particles Area in LD were quantified with the ImageJ ‘‘analyze particles’’ function in thresholded images, with size (square µm) settings from 0.1 to 100 and circularity from 0 to 1. To determine Red C12 % area per cell, Red C12 paritcles in LD area was normalized with Cell area calculated with Cellpose generalist algorithm for cell segmentation.

LD (BODIPY 493/503) or Lysosome (LysoTracker™ Red DND-99) particle area was quantified with the ImageJ ‘‘analyze particles’’ function in thresholded images, with size (square µm) settings from 0.1 to 100 and circularity from 0 to 1. To determine LD or Lysotracker % area per cell, LD or lysosomal particles were normalized with Cell area. Cell area was automatically generated using Cellpose generalist algorithm for cell segmentation. Data were expressed as means ± SEM. Statistical analysis among groups was performed using Student’s t test.

**SeaHorse XF-96 metabolic flux analysis**

For Seahorse metabolic flux experiments, oxygen consumption rates and extracellular acidification rates were measured using a 96-well Seahorse Bioanalyzer XF 96 according to the manufacturer’s instructions (Agilent Technologies).

*XF Cell Mito Stress Test*

Fibroblast cells were seeded (18,000 cells per well) into 96-well plates the previous day to the assay. Prior to assay, starving condition cells were switched to starving medium for 6h before analysis. Analysis was performed according to the manufacturer´s instructions. 1h prior to analysis media was replaced with 175uL Mito assay medium and incubated for 1h at 37ºC without CO2. During the experiments inhibitors were sequentially injected: Oligomycin [2uM], FCCP [2uM], Rotenone [0.5uM] and Antimycin [0.5uM]. Then OCR was automatically calculated by the Seahorse XF-24 analyser (Seahorse Bioscience, CA, USA).

ATP production calculated as the difference between the last rate measurement before Oligomycin injection and the minimum rate measurement after Oligomycin injection. mitochondrial maximal respiration calculated as the di_erence between the maximum rate measurement after FCCP injection and the non-mitochondrial respiration. mitochondrial spare respiratory capacity calculated as the difference between the maximal and basal respiration.

Mito assay medium: 8.7g/L MEM (61100-087, ThermoFisher scientific), 1mM pyruvate (GIBCO), 2mM glutamine and 10mM glucose, pH=7.4.4

*The XF Palmitate Oxidation Stress Test*

At 48 h before the experiment, cells were cultured on XF-96 plates at a density of 18.000 cells/well. At 24 h before metabolic flux analysis, the culture medium was replaced with 100 µL of substrate-limited medium. Before the analysis, the medium was replaced with 135 µL FAO Assay Medium and incubated at 37°C in a non-CO2 incubator for 1h. Just prior to assay cells were treated with palmitate-BSA (200 µM) or BSA (34 µM), and during the experiment inhibitors were sequentially injected: Etomoxir (40 µM), Olygomycin (2uM), FCCP (2uM), antimycin A (0.5 µM) and rotenone (0.5 µM). Then OCR was automatically calculated by the Seahorse XF-24 analyzer (Seahorse Bioscience, CA, USA).

Basal FFA Oxidation calculated as the difference between the second rate measurement after Etomoxir injection and the last rate measurement before Etomoxir injection. ATP production calculated as the diference between the last rate measurement before Oligomycin injection and the minimum rate measurement after Oligomycin injection. mitochondrial maximal respiration calculated as the di_erence between the maximum rate measurement after FCCP injection and the non-mitochondrial respiration.

Substrate-Limited Medium: Seahorse XF DMEM (103575-100, Agilent), 0.5 mM D-Glucose (141341, ITW reagents), 1mM GlutaMAX (35050038, Gibco), 05 mM L-Carnitine (C0158, Sigma Aldrich ), 1% FBS (10270-106, Gibco), pH=7.4.

FAO assay buffer: Seahorse XF DMEM ((103575-100, Agilent), 0.5 mM L-Carnitine (C0158, Sigma Aldrich ), pH=7.4.

**Transmission electron microscopy TEM**

Transmission electron microscopy After 72 h transfection, the MCF-7 GFP-LC3B cells were fixed in 3% v/v glutaraldehyde in 0.1 M sodium phosphate buffer (pH 7.4) for 10min at 37ºC and 2h at room temperature (RT). The samples were washed five times in 0.1 M sodium phosphate buffer (pH 7.4). The fixed cells were delivered to Unidad de Neurobiología Comparada of the University of Valencia (group of Dr. García-Verdugo). Once there, samples were treated with 2% Osmium tetroxide in PBS (pH 7.2, Ca2 + and Mg2 + free, Gibco) for 2 hours at room temperature, washed, dehydrated with increasing ethanol solutions and dyed with 2% Acetate Uranyl in 70% ethanol. The dehydrated samples were embedded in Araldite. Semi-thin sections (1.5 μm) were cutted and stained with 1% Toluidine blue solution. After this, ultra-thin sections (70nm) were done to be examined under transmission electron microscopy Tecnai-Spirit coupled to Morada TEM CCD (Soft Imaging System) camera. Lysosomes, Autophagosomes, Fingerprint structures, lipid droplet and mitochondria area were quantified in 10–15 cells per sample cristae measurement as (10.3390/cells10092177).
