## Supplementary Figures for "Lipid Dysregulation Unveil the Intricate Interplay of Lysosomal and Mitochondrial Changes in Frontotemporal Dementia with GRN Haploinsufficiency"

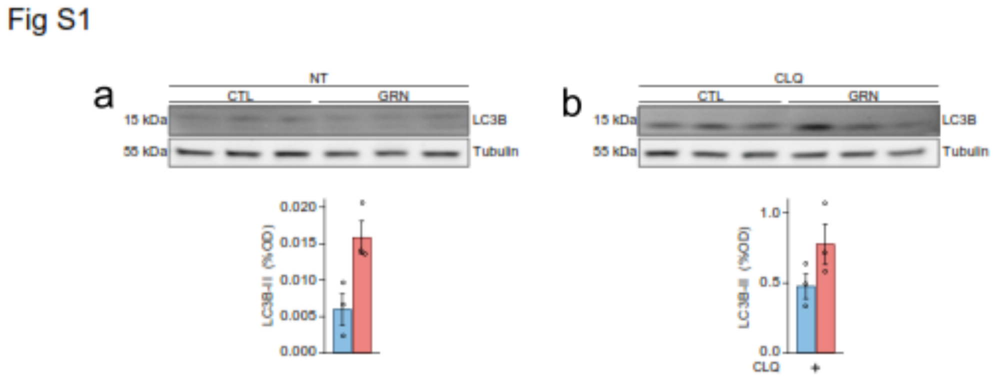


**Figure S1. Extension of Figure 3.**

A. Protein Levels of LC3B-II Assessed by Western Blotting in Primary Human Fibroblasts. Western blotting was performed to assess the protein levels of LC3B-II, and quantification are shown as bar plot graphs.

B. Protein Levels of LC3B-II Assessed by Western Blotting in Primary Human Fibroblasts. Primary human fibroblasts were treated with CLQ at 30 μM for 5 hours. Western blotting was performed to assess the protein levels of LC3B-II, and quantification are shown as bar plot graphs.

Healthy control fibroblasts(CTL), FTD-GRN patient fibroblast (GRN). Data are presented as means ± SEMs ($n = 3$). Statistical significance denoted by $p > 0.05$, determined using Student's t test.

Fig S2

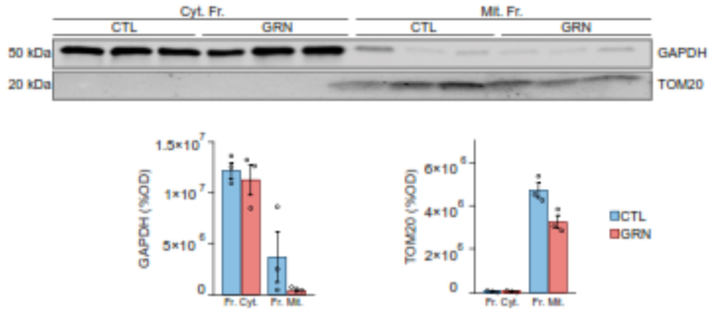


**Figure S2. Cytoplasmic and Mitochondria fractionation.**

A. Protein Levels of GAPDH and TOM20 Assessed by Western Blotting in Primary Human Fibroblasts. Western blotting was performed to assess the protein levels of GAPDH and TOM20, and quantification are shown as bar plot graphs.

Healthy control fibroblasts(CTL), FTD-GRN patient fibroblast (GRN). Data are presented as means ± SEMs (n = 3). Statistical significance denoted by $p > 0.05$, determined using Student's t test.

Fig S3

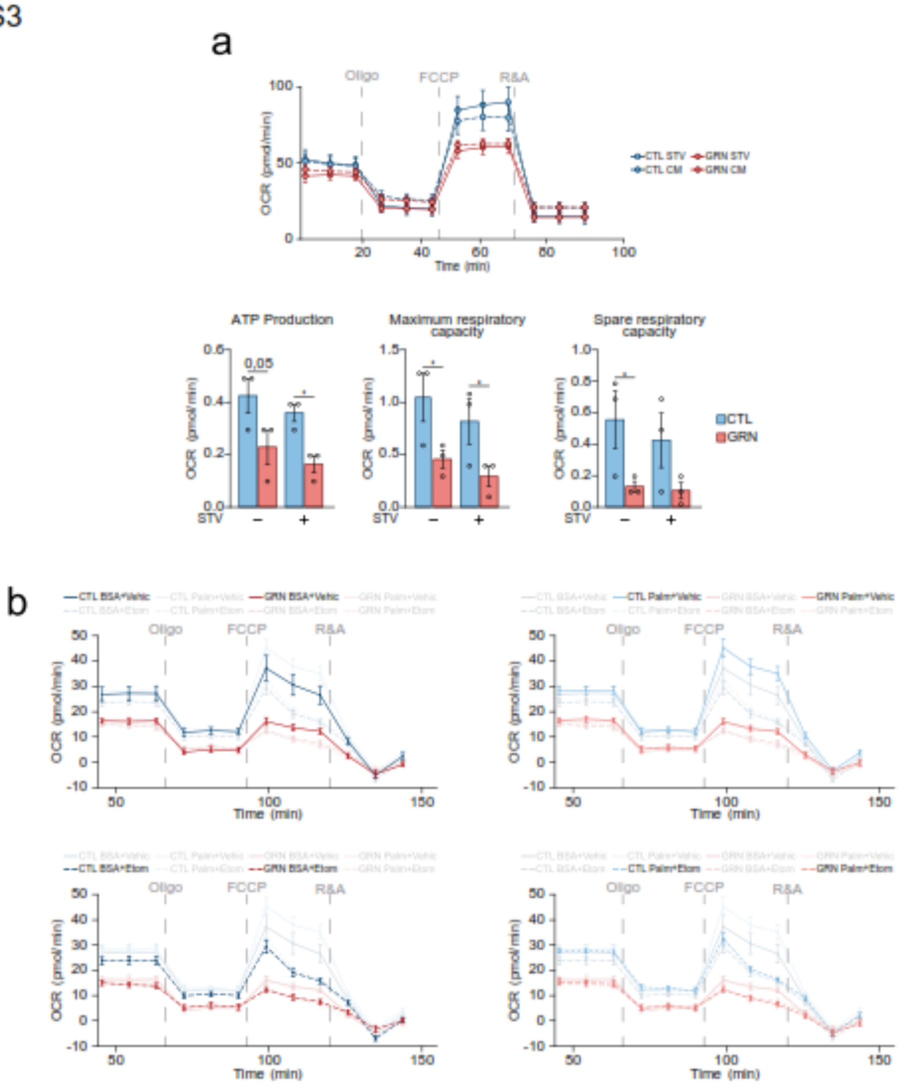


**Figure S3. Extension of Figure 4.**

A. Fig 4C. Seahorse Assay depicting Mitochondrial Oxygen Consumption in Primary Human Fibroblasts. Primary human fibroblasts were cultured with or without a 6-hour STV pre-assay period.

B. Fig 4G in detail.

Healthy control fibroblasts(CTL), FTD-GRN patient fibroblast (GRN). Data are presented as means ± SEMs (n = 3).

Fig S4

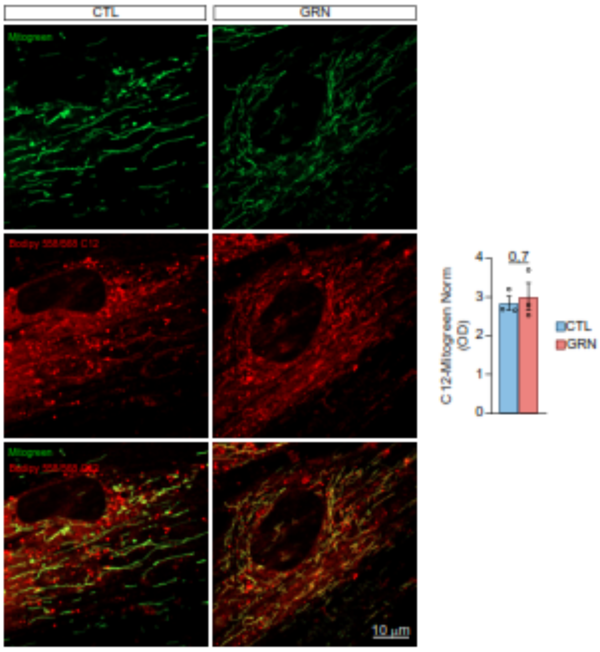


**Figure S4. FA inside Mitochondria.**

A. Representative Images of Mitogreen (green) and TOM20 (red) Staining in Primary Human Fibroblasts. Scale bar = 10 µm.

Healthy control fibroblasts (CTL), FTD-GRN patient fibroblast (GRN). Data are presented as means ± SEMs (n = 3).

Fig S5

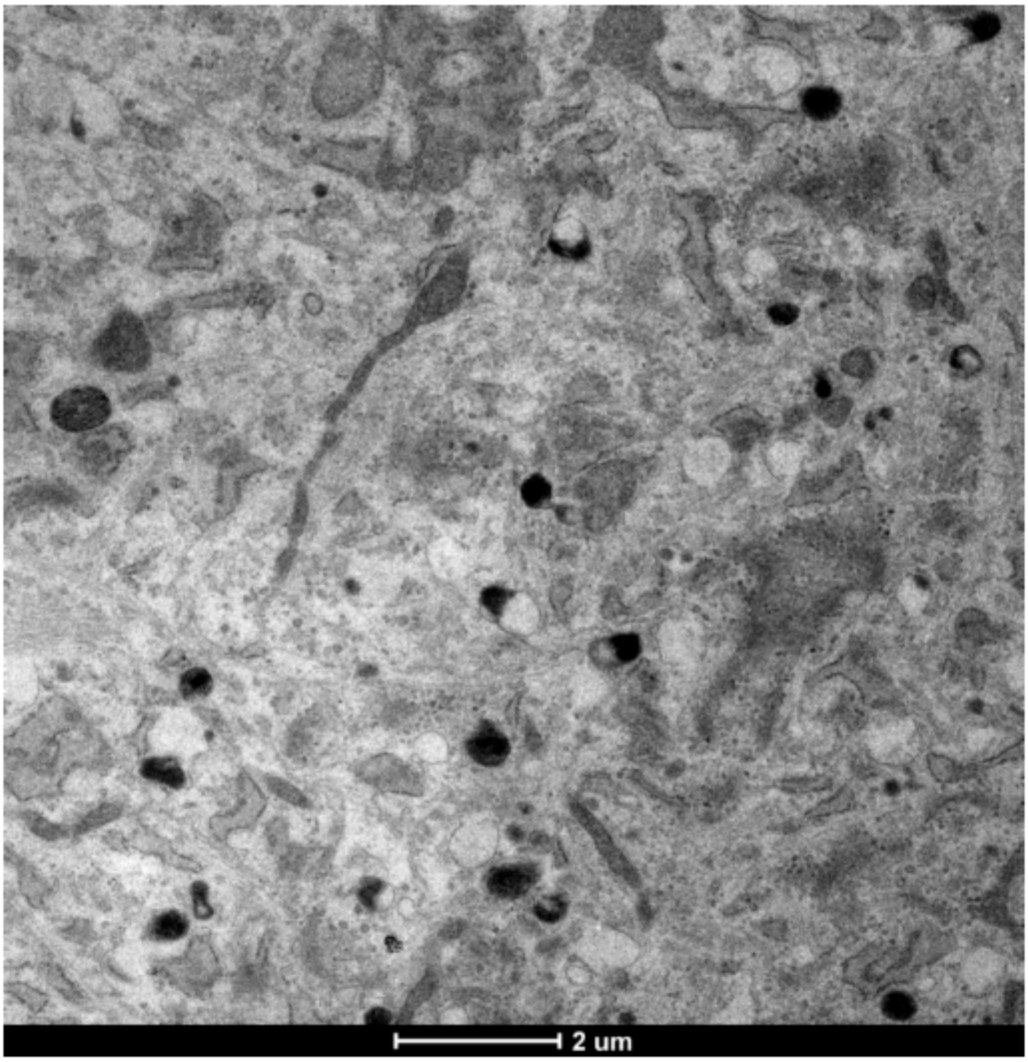


**Figure S5. TEM image in detail.**

TEM Images of healthy control Primary Human Fibroblasts 03. Scale bar = 2μm.

Fig S6

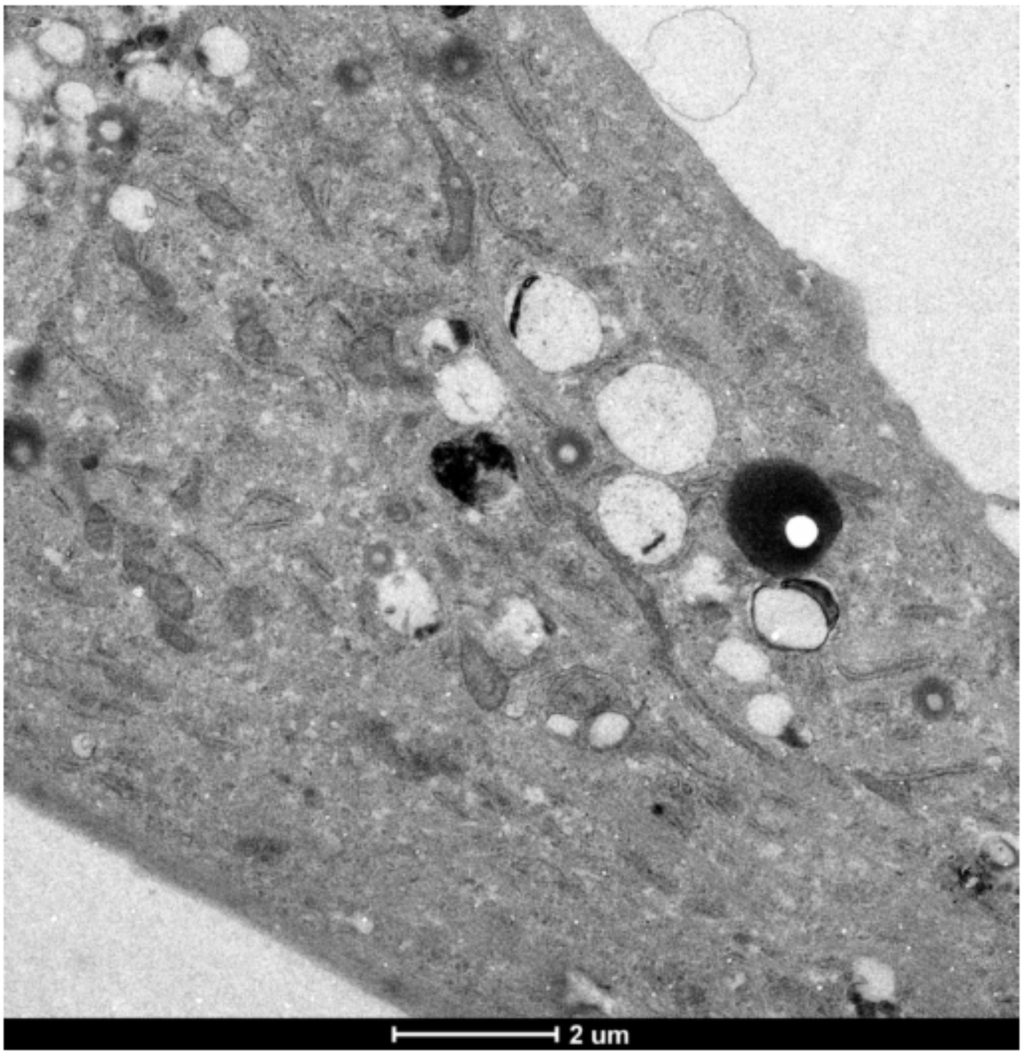


**Figure S6. TEM image in detail.**

TEM Images of FTD-GRN patient Primary Human Fibroblast 04. Scale bar = 2µm.

Fig S7

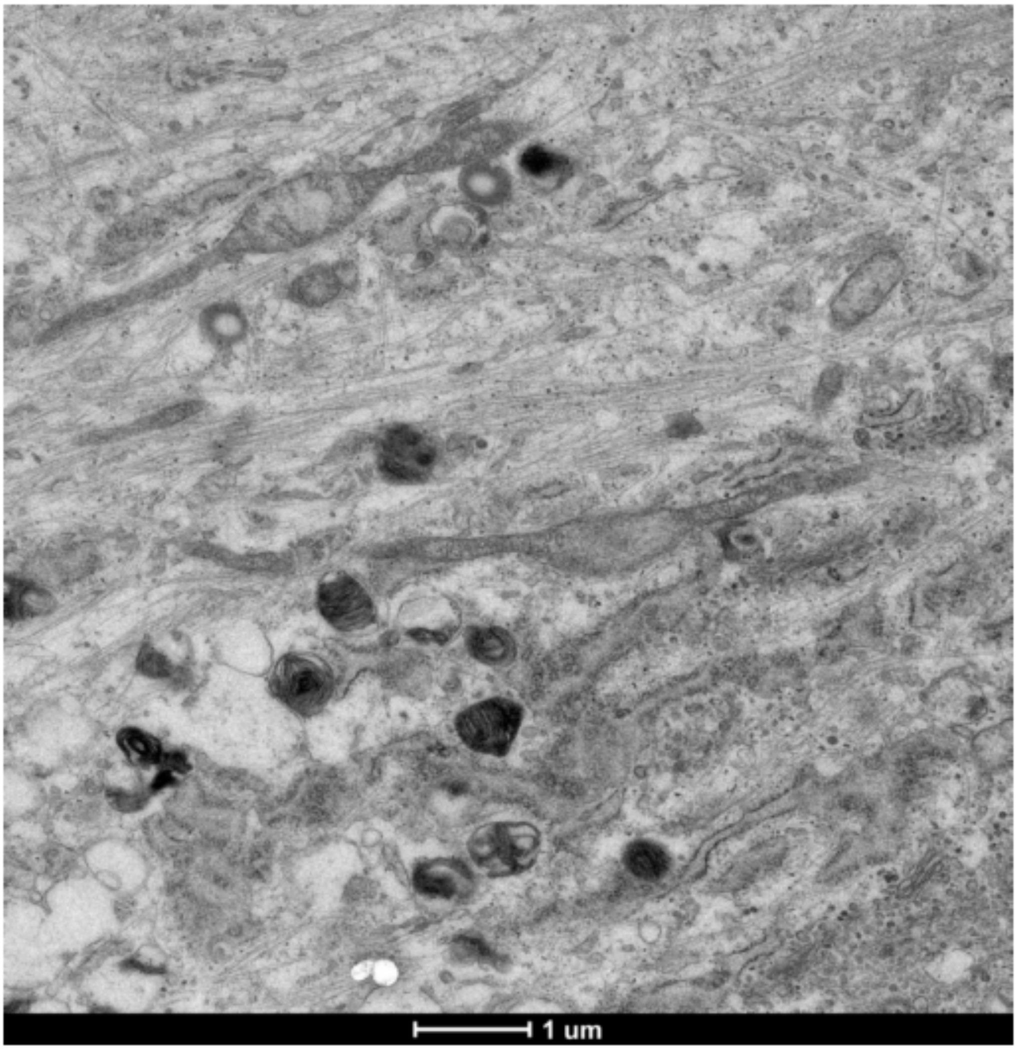


**Figure S7. TEM image in detail.**

TEM Images of FTD-GRN patient Primary Human Fibroblast 05. Scale bar = 1μm

Fig S8

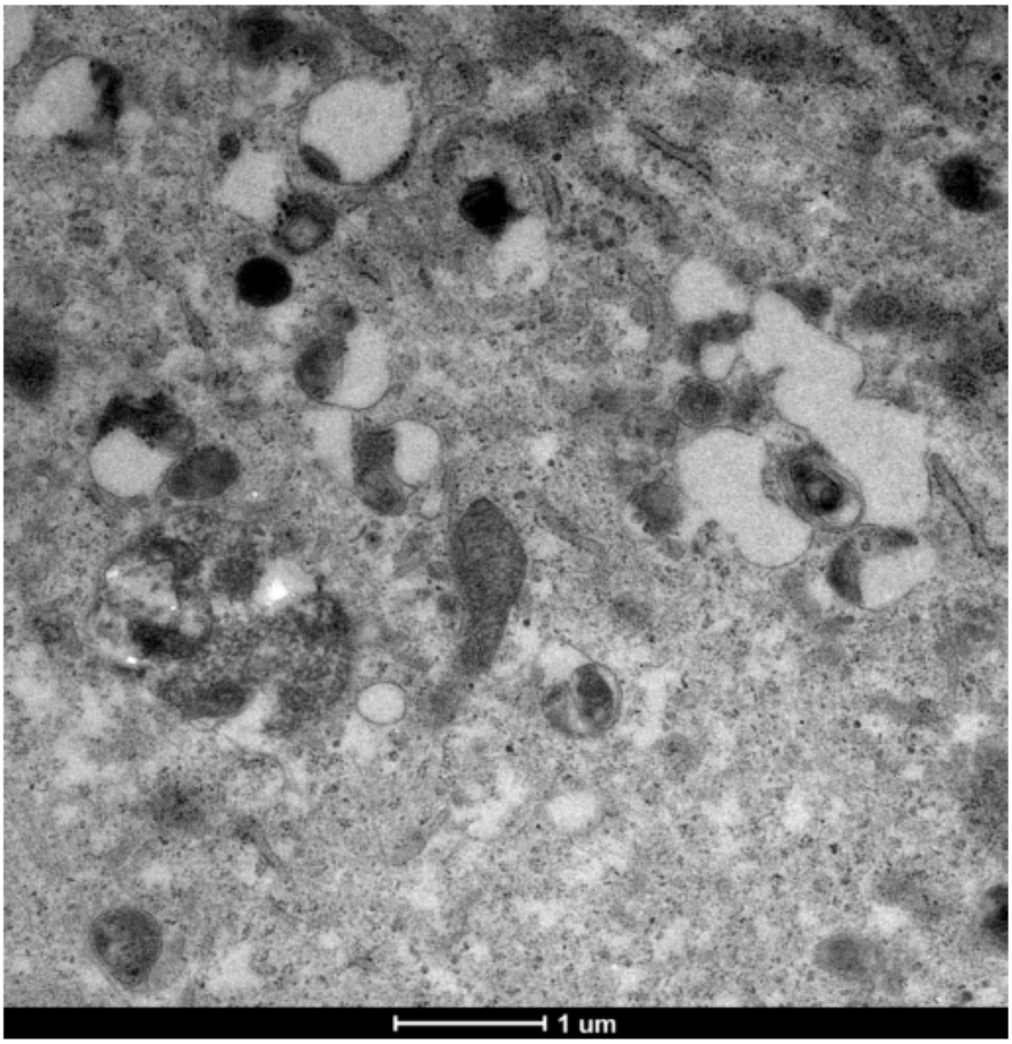


**Figure S8. TEM image in detail.**

TEM Images of healthy control Primary Human Fibroblasts 02. Scale bar = 1μm

Fig S9

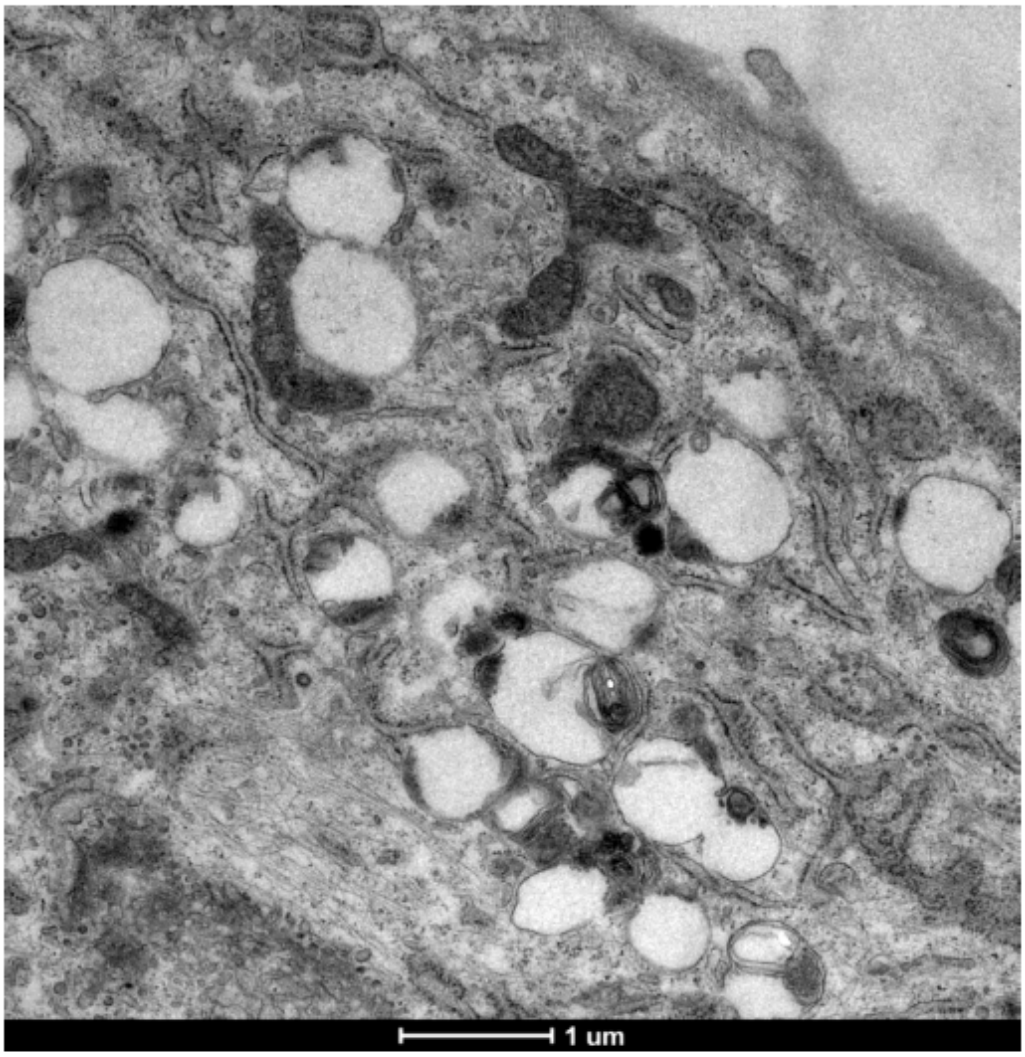


**Figure S9. TEM image in detail.**

TEM Images of FTD-GRN patient Primary Human Fibroblast 05. Scale bar = 1μm

Fig S10

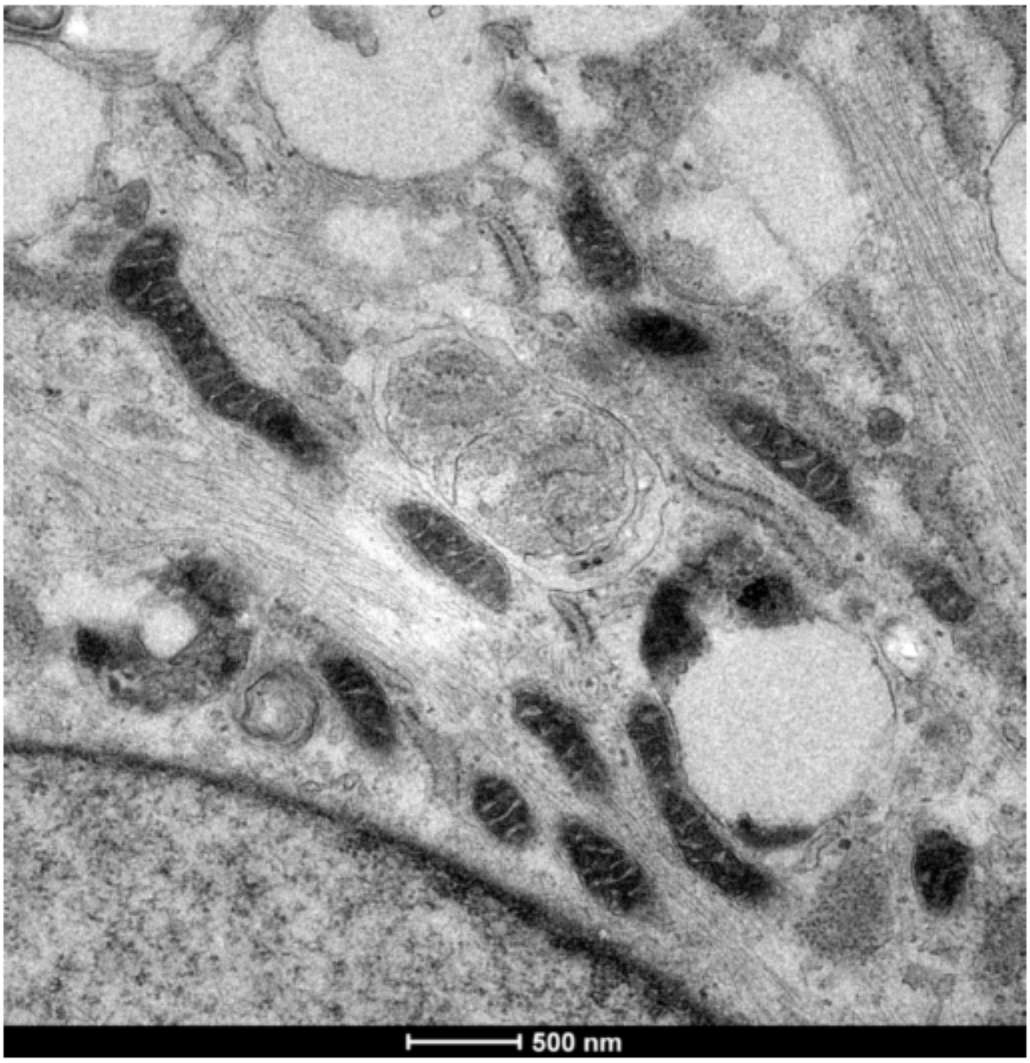


**Figure S10. TEM image in detail.**

TEM Images of healthy control Primary Human Fibroblasts 03. Scale bar = 500nm

Fig S11

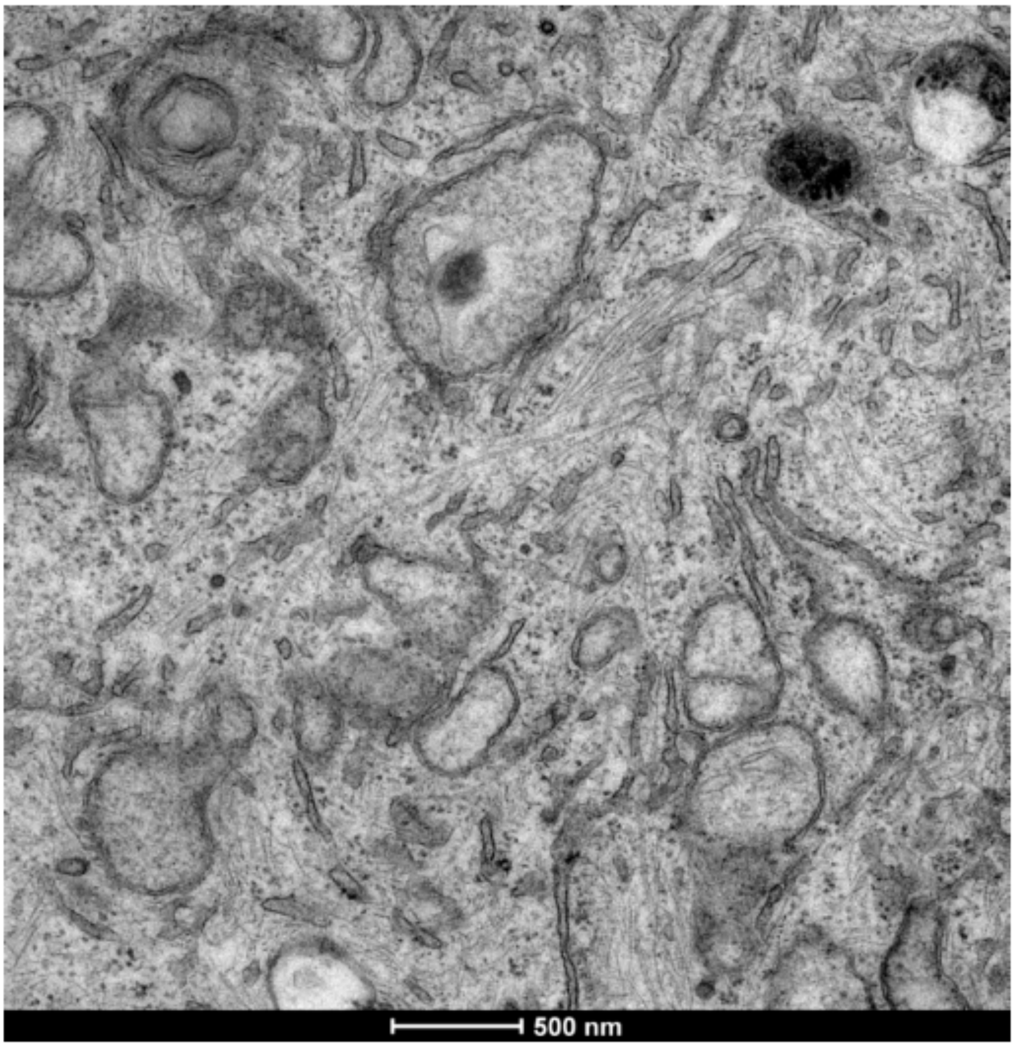


**Figure S11. TEM image in detail.**

TEM Images of FTD-GRN patient Primary Human Fibroblast 05. Scale bar = 500nm

Fig S12

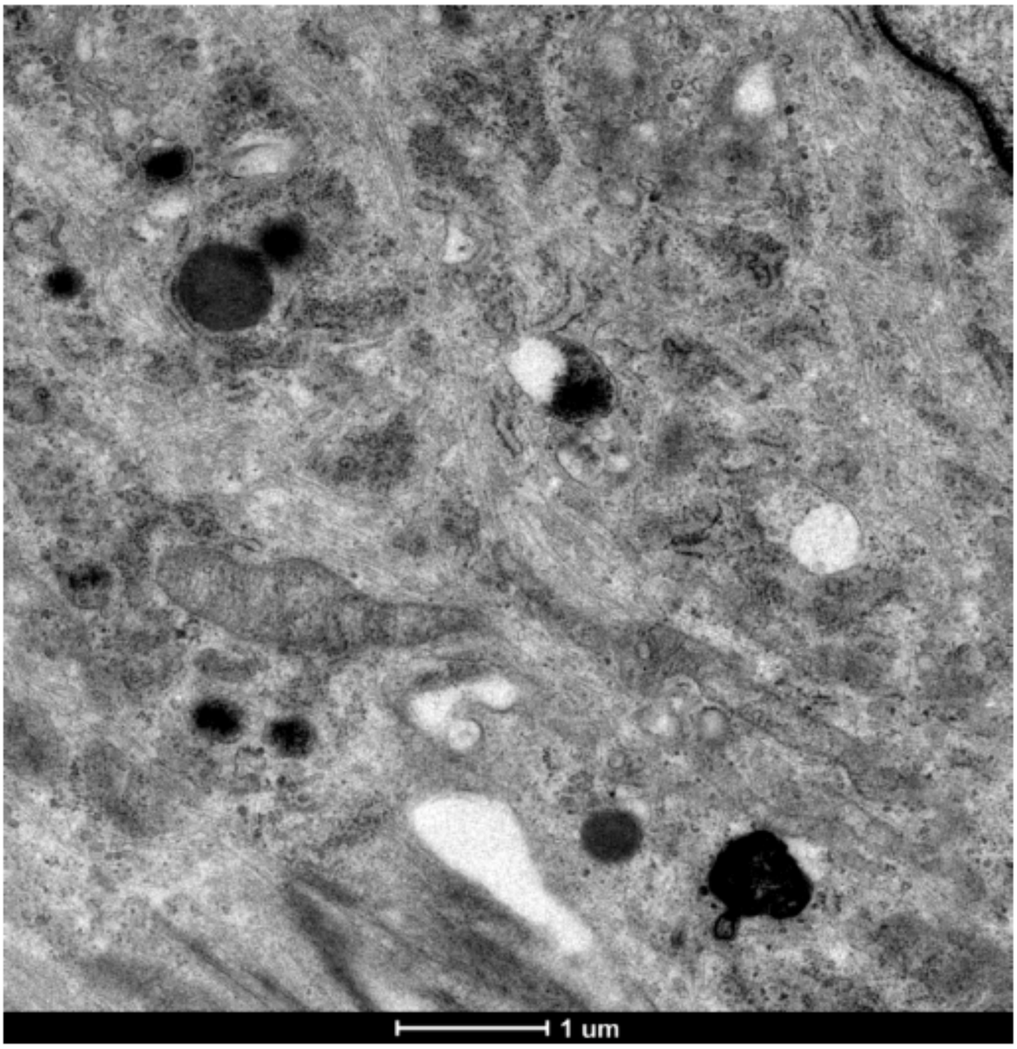


**Figure S12. TEM image in detail.**

TEM Images of healthy control Primary Human Fibroblasts 01. Scale bar = 1µm

Fig S13

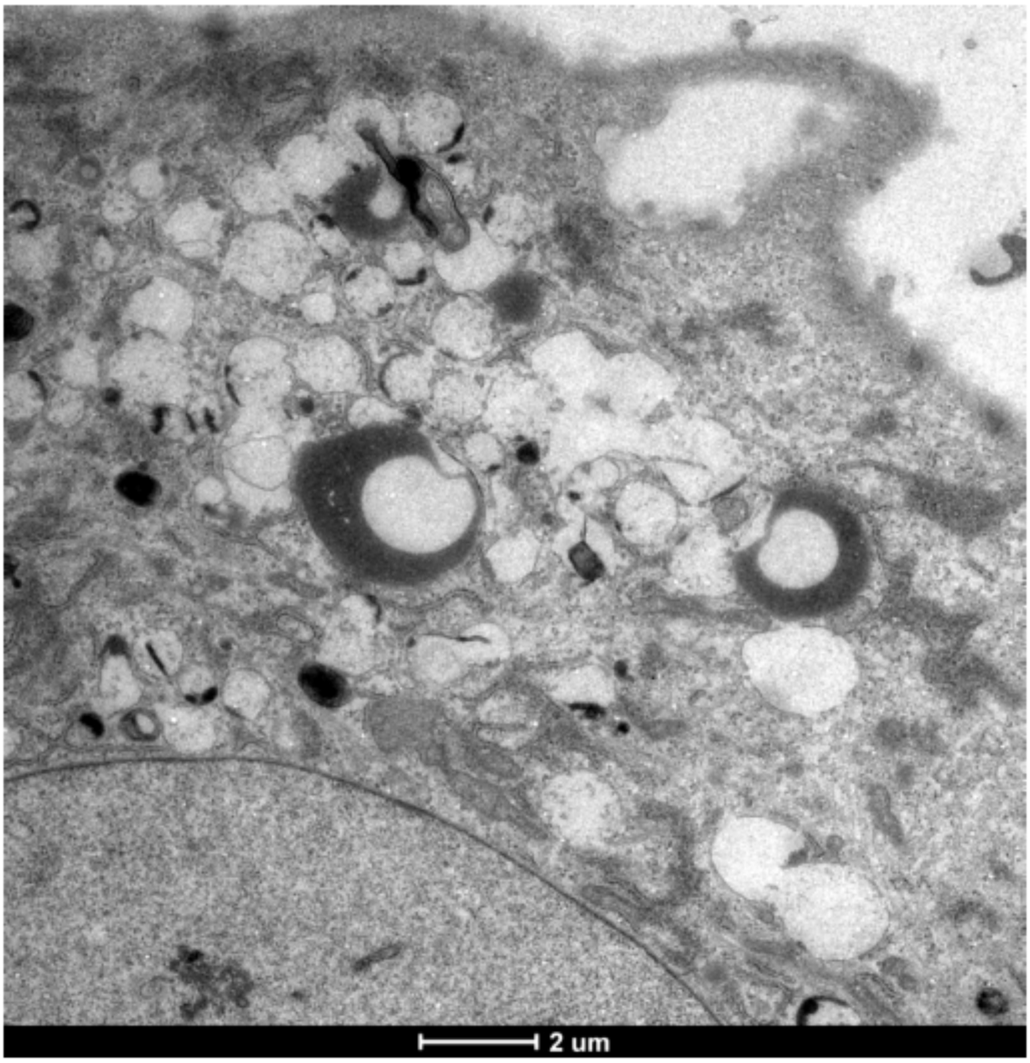


**Figure S13. TEM image in detail.**

TEM Images of FTD-GRN patient Primary Human Fibroblast 04. Scale bar = 2µm
